# Transcriptional responses of brain endothelium to *Plasmodium falciparum* patient-derived isolates *in vitro*

**DOI:** 10.1101/2024.01.11.575088

**Authors:** Caroline Askonas, Janet Storm, Grazia Camarda, Alister Craig, Arnab Pain

**Affiliations:** Pathogen Genomics Laboratory, Bioscience Program, Biological and Environmental Sciences and Engineering (BESE), Division, King Abdullah University of Science and Technology, Thuwal, KSA; Tropical Disease Biology, Liverpool School of Tropical Medicine, Pembroke Place, Liverpool L3 5QA, UK

**Keywords:** cerebral malaria, cytoadherence, Plasmodium falciparum, HBMEC, endothelium, gene expression

## Abstract

A hallmark of cerebral malaria (CM) is sequestration of *Plasmodium falciparum* infected erythrocytes (IE) within the brain microvasculature. Binding of IE to endothelium reduces microvascular flow and combined with an inflammatory response perturbs endothelial barrier function, resulting in breakdown of the blood-brain barrier (BBB). Cytoadherence leads to activation of the endothelium and alters a range of cell processes affecting signalling pathways, receptor expression, coagulation, and disruption of BBB integrity. Here, we investigated whether CM derived parasites elicit differential effects on human brain microvascular endothelial cells (HBMECs), as compared to uncomplicated malaria (UM) derived parasites. Patient-derived IE from UM and CM clinical cases, as well as non-binding skeleton-binding protein 1 knockout parasites, were overlaid onto TNF-activated HBMECs. Gene expression analysis of endothelial responses was performed using probe-based assays of a panel of genes involved in inflammation, apoptosis, endothelial barrier function, and prostacyclin synthesis pathway. We observed a significant effect on endothelial transcriptional responses in the presence of IE, yet there was no significant correlation between HBMEC responses and type of clinical syndrome (UM or CM). Further, there was no correlation between HBMEC gene expression and both binding itself and level of IE binding to HBMEC, as we detected the same change in endothelial responses when employing both binding and non-binding parasites. Our results suggest that interaction of IE with endothelial cells in this co-culture model induce some endothelial responses that are independent of clinical origin and independent of the expression of the major variant antigen *Pf*EMP1 on the IE surface.

## Introduction

In spite of global efforts to reduce mortality and morbidity of malaria, an estimated 249 million cases and 608,000 deaths were reported in 2022, with the majority occurring in children under the age of five ^1^. The majority of symptomatic infections result in mild febrile cases of uncomplicated malaria (UM), as severe disease only occurs in 1-2% of all cases ^2^. Out of the eight *Plasmodium spp* that have been reported to cause either direct or zoonotic infection in humans, *P. falciparum* is considered a major driver of severe disease and mortality ^3,4^. Cerebral malaria (CM) is a severe neurological complication of malaria infection that is a major cause of death, with mortality estimated between 15-25% and clinical manifestations including unarousable coma and seizures ^5,6^. Mortality in paediatric CM is associated with brain swelling, which is considered to occur as a result of increased permeability of the blood-brain barrier (BBB) due to loss of BBB function, leading to vasogenic oedema ^6,7^. Continued study of the pathophysiology of CM is required to develop targeted treatment therapies.

It is evident that CM is a complex, multi-component disease requiring a multifactorial view of pathogenesis that considers the heterogeneous clinical manifestations of CM ^8,9^. Sequestration of *P. falciparum*-infected erythrocytes (IE) to brain microvasculature is a hallmark of CM. Cytoadhesion of IE to the endothelium is mediated by *Plasmodium falciparum* erythrocyte membrane protein 1 (PfEMP1), a variable surface protein that is expressed by IE and contains an extracellular binding region ^10^. This polymorphic protein is encoded by a group of approximately 60 *var* genes with monoallelic expression, and the composition of the extracellular region confers binding to various host receptors ^11^. Several vascular surface proteins, in particular CD36, EPCR, and ICAM-1, may serve as IE adhesion receptors, and expression of these endothelial receptors varies between vascular locations and is modulated in part by cytokines, such as TNF and IFN ^9,10,12,13^. EPCR is associated with severe malaria, as PfEMP1s containing EPCR binding domains have been linked to brain swelling in paediatric CM ^14–17^. While ICAM-1 mediates cytoadhesion related to sequestration in the brain ^13–15^, its direct role in the pathology of CM is less clear. However, research has demonstrated that a subset of EPCR-binding parasites has the ability to bind ICAM-1and that expression of these dual binders has been associated with CM in patients ^15,18,19^. Cytoadherence alters a range of endothelial cell processes and leads to the activation of endothelium as well as reducing microvascular flow. Accumulation of IE causes vessel occlusion and generates microenvironments where both parasite factors and soluble erythrocytic content accumulate after IE rupture ^8,20^. Ultimately, the contributions of microvascular obstruction and endothelial dysfunction lead to the disruption and breakdown of the BBB^5,8,9,17^.

Although IE derived from malaria patients have exhibited different binding phenotypes, most studies employ well characterized laboratory strains to investigate cytoadherence and interaction of different PfEMP1 variants to brain endothelial cells ^19,21–24^. Separate investigations have highlighted the divergent effect of both TNF stimulation and IE cytoadhesion on endothelial transcriptional responses and the pathways they induce ^23,25^. IE isolated from peripheral blood of UM and CM patients have been used to study cytoadherence to human brain microvascular endothelial cells (HBMEC). It was found that IE isolated from CM patients exhibit differential binding capacities to TNF-activated brain endothelium *in vitro*, as they were associated with increased adhesion ^26^. Further study employing patient-derived isolates is needed to elucidate the impact of parasite cytoadhesion on the brain microvasculature in association with different disease outcomes.

The aim of this study is to investigate whether CM-derived IE elicit differential effects on HBMEC as compared to UM-derived isolates. An *in vitro* co-culture model of cytoadherence was employed wherein patient-derived IE isolates were overlayed onto TNF-activated HBMEC and the relative gene expression of HBMEC after incubation with either RBC or IE for a panel of genes was assessed. A study utilizing two *P. falciparum* lab strains with different binding capacities to HBMEC found that different PfEMP1-expressing variants induced divergent endothelial transcriptional responses during cytoadherence ^25^. Differentially expressed genes (DEGs) were identified in HBMEC in pathways involved in the pathology of severe malaria, such as inflammation, apoptosis, and barrier integrity. From this list of DEGs, a panel of genes was compiled, and gene expression was investigated by qPCR using the Fluidigm DELTAgene assays system. First, the relative gene expression of HBMEC after exposure to control RBC or IE for combined UM and CM-derived isolates was compared, followed by comparing the expression data between clinical type. We observed variation in endothelial transcriptional responses to individual patient-derived IE as compared to RBC, but these responses did not show a significant differential effect between the UM or CM-derived isolates. It was assessed whether the HBMEC relative gene expression after co-culture was correlated with the binding avidity of the parasite isolates, but a significant relationship was not observed. Our data confirm that IE have a profound effect on endothelial transcription that is specific to the presence of the *P. falciparum* parasites in the erythrocyte. However, we did not observe effects that were associated with disease severity with the panel of genes used in this study.

## Materials and Methods

### Culture of Endothelial Cells

Primary HBMEC were obtained from Cell Systems, USA (ACBRI 376) and cultured in Endothelial Cell Growth Medium 2 (EGM2) media (C-22111, Promocell) supplemented with 2% foetal calf serum (FCS), 5 ng/mL epidermal growth factor (EGF), 10 ng/mL basic fibroblast growth factor, 20 ng/mL Long R3 IGF-1, 0.5 ng/mL VEGF 165, 1 µg/mL ascorbic acid, 22.3 µg/mL heparin, and 0.2 µg/mL hydrocortisone from the supplement pack (C-39211, Promocell) in a 5% CO2 incubator at 37°C. For endothelial cells to adhere to the surface, flasks were coated with attachment factor (10308363, Gibco). Cells were passaged using Accutase solution (A6964, Sigma Aldrich), following manufacturer instructions. The receptor expression of HBMEC was characterised by flow cytometry as described by Storm et al ^26^.

### Culture of *P. falciparum* Parasites

Patient-derived *P. falciparum* parasites were isolated from peripheral blood from paediatric malaria cases recruited at the Queen Elizabeth Central Hospital, Blantyre, Malawi with ethical approval from the College of Medicine, University of Malawi and LSTM, as described in Storm et al ^26^. Four UM and four CM patient-derived isolates were selected based on their differential cytoadherence profile to HBMEC and human dermal microvascular endothelial cells and their ability to establish in continuous culture (Figure 3). After isolation from blood, limited IE could be cryopreserved, especially for CM isolates, and only a maximum of 150 µl IE could be frozen. Establishing the isolates as a continuous culture, expanding the cultures to generate more stabilates, and subsequently have 65 ml of culture volume at 2% HCT for the co-cultures requires a considerable amount of time. Therefore, co-cultures were performed after at least 21 days in culture (DIC). The total DIC for the patient-derived parasite isolates for co-culture 1 and 2, respectively, were UM1: 24 and 43, UM2: 21 and 32, UM3: 31 and 42, UM4: 24 and 25, CM1: 29 and 29, CM2: 35 and 44, CM3: 49 and 59, and CM4: 29 and 41. The skeleton-binding protein knockout (SBP1-KO) lab strain was obtained from A.G. Maier and cultured in the presence of 4 nM WR99210 ^27^.

The IE were cultivated at 2% haematocrit in O+ human erythrocytes and grown at 37°C in multiple 25cm^2^ tissue culture flasks (T25) filled with a gas mixture of 96% nitrogen, 3% carbon dioxide, and 1% oxygen. The duration of parasite culturing was isolate dependent, as different lengths of continuous culture were required to generate enough material to perform the experiments, ranging from 21-60 DIC. IE were maintained in complete RPMI 1640 medium with 11 mM glucose and 0.2% sodium bicarbonate (R-0883, Sigma) supplemented with 5% human serum, 0.25% Albumax II (11021029, ThermoFisher Scientific), 30 mM of HEPES (15630122, Gibco), 2 mM L-glutamine (G-7513, Sigma), 25 ng/mL gentamicin sulphate solution (G-1271, Sigma), and 0.11 mM hypoxanthine (H9636, Sigma) pH 7.4. Parasite growth was monitored using Giemsa staining (1092040500, Merck), and IE were passaged to maintain 2-5% parasitaemia. Mixed-staged cultures were synchronized for ring stages using a 5% D-sorbitol solution (S3889, Sigma), and IE at trophozoite stage were used for the *in vitro* co-culture studies with HBMEC. EC cultures, parasite lines, culture media, and washed RBCs were monitored for mycoplasma contamination using MycoBlue™ Mycoplasma Detector (Vazyme, D101-02).

### Co-culture of HBMEC and IE

HBMEC at passage 5-8 were cultured until 90% confluency in T25 flasks. One day prior to co-culture, HBMEC were seeded in 12-well plates at a density of 50,000 cells/cm^2^ (for 3.8 cm^2^ wells, this is equivalent to total of 190,000 cells) in the morning and stimulated overnight with 10 ng/mL TNF (PHC3015, Invitrogen). The RBC control was prepared by culturing a 2% haematocrit suspension in RPMI medium overnight. On the day of the co-culture, the supernatant was removed from the HBMEC and replaced with EGM2 media without heparin and hydrocortisone (EGM2min media) 2 hours prior to co-culture. IE were enriched for mature stages by layering onto a 0.7% gelatine solution and incubation at 37°C for 45-60 minutes and after enrichment, the haematocrit and parasitaemia were calculated and IE suspensions were prepared to an average parasitaemia of 30% (parasitaemia ranged from 20-40%, depending on enrichment properties of the isolates) at 1% haematocrit in EGM2min media.

The RBC control was also subjected to gelatine treatment and a suspension at 1% haematocrit in EGM2min media was prepared. EGM2min medium was used to maintain healthy HBMEC and did not affect parasite viability in the 6-hour co-culture, but parasite development seemed delayed, as assessed by microscopy. Medium was removed from the HBMEC, and 700 µl IE or RBC suspension were overlayed onto HBMEC for 6 hours in a 5% CO2 incubator at 37°C, as well as EGM2min medium as an additional control. 700 µL of 30% parasitaemia at 1% haematocrit is equivalent to 2.1×10^7^ IE and addition of the IE suspension to HBMEC is equivalent to a ratio of 110 IE/HBMEC. A 0-hour control with EGM2min medium was used to compare with the 6-hour time points. Control samples were generated by overnight incubation of HBMEC with either 10 ng/mL of TNF, or 1 ng/mL IL-1 beta (579402, BioLegend) and compared to EGM2min medium, for which samples were collected directly after stimulation.

At each time point, the co-culture medium was removed, and the HBMEC were washed once with EGM2min medium to remove unbound IE or uninfected RBC. Cells were directly lysed in the well using 350 µL of lysing solution (1% 2-mercaptoethanol in Buffer RLT; RNeasy Mini Kit, 74106, Qiagen) by pipetting over the well. The lysates were collected into a 1.5mL sterile tube, vortexed, and stored at −80°C until RNA extraction. Two independent co-culture experiments were performed for each patient isolate and SBP1-KO.

In parallel to the co-culture experiments, activation of HBMEC after overnight incubation with TNF was assessed. One day prior to co-culture, HBMEC were plated onto 24-well plates at a density of 50,000 cells/cm^2^. The HBMEC were incubated overnight with either media alone or media containing 10 ng/mL of TNF. Cells were detached with Accutase, collected in PBS + 1% FBS and stained for ICAM-1 expression on the endothelial cell surface using APC anti-human CD54 antibody (353111, Biolegend). Flow cytometry analysis was performed and confirmed that ICAM-1 expression was upregulated following overnight stimulation with TNF in each of the co-culture experiments.

### RNA Extraction and Quality Assessment

Frozen HBMEC lysates were placed in a 37°C water bath until thawed and salts dissolved. Total RNA was purified using the RNeasy Mini Kit (74106, Qiagen) following the manufacturer’s instructions using steps 4-10 of the “Purification of Total RNA from Animal Cells using Spin Technology” protocol. RNA was eluted in 50 µL of RNase-free water.

The RNA concentration of each sample was quantified using a Qubit fluorometer and the RNA High Sensitivity assays (Q32852, Invitrogen), following manufacturer instructions. The quality of the RNA samples was assessed using an Agilent 2100 Bioanalyzer (RNA 6000 Nano Kit, 5067-1511, Agilent technologies) following manufacturer instructions. The RIN for all samples was between 9.4-10, indicating high quality RNA.

### Gene Expression Assays using the Fluidigm System and Calculations

Gene expression qPCR assays were conducted on the extracted RNA for a panel of 49 genes that include pathways pertaining to inflammation, apoptosis, endothelial barrier function, and prostacyclin synthesis (listed in Supplementary Table 1). In short, cDNA was prepared with reverse transcription before pre-amplification (13-cycles) and clean-up of the reactions using Exonuclease 1, following the manufacturer protocols (100-6472 B1, PN 100-5875 C1) and reagents (Fluidigm,100-6297; Fluidigm, 100-5580; New England Biolabs, M0293S; TEKnova, 10221). Real-time PCR data was collected using the Biomark HD system with a 96.96 IFC chip using Fluidigm Delta-Gene Assays on the pre-amplified cDNA. The manufacturing protocol was followed for use of the Biomark HD system (PN 100-9792 A1) using the reagents specified for preparing samples (Biorad, 172-5211; Fluidigm, 100-7609)) and assays (Fluidigm 100-7611; TEKnova, 10221). Technical replicates were performed for each sample, and Ct values were obtained as the throughput of the real-time data. The primers for the gene panel were proprietarily designed, prepared, and tested by Fluidigm.

Relative gene expression was calculated using the 2^−ΔΔCt^ method as previously described ^28^ using GAPDH as the endogenous control (ΔCt) and the 6-hour EGM2min media-only sample as the normalization controls (ΔΔCt). In this way, the exponential fold changes (FCs, 2^−ΔΔCt^) in the target genes are calculated as normalized to the GAPDH internal control and relative to the expression of the HBMEC-media only at the 6-hour time point. For each target gene, the gene expression of HBMEC incubated with EGM2min media at 6-hours is equal to 1 (baseline). Therefore, an FC>1 indicates increased relative gene expression and an FC <1 indicates decreased relative gene expression for the experimental samples. FCs were calculated for each technical triplicate of each sample for each experiment. To compare relative gene expression of HBMEC after incubation with various groups (e.g., RBCs, UM-derived parasite isolates, CM-derived parasites isolates, or combined IE), the mean, standard error of the mean (SEM), and confidence intervals for each group were calculated from the technical triplicates of each sample for each independent experiment. Either two-tailed, unpaired T-tests or Welch’s T-test were then performed to compare the relative gene expression between various corresponding groups. Additionally, a paired analysis was performed wherein the HBMEC-IE relative gene expression for each parasite isolate was compared to the corresponding HBMEC-RBC relative gene expression, for both experiments. For this analysis, the target FCs were calculated as normalized to GAPDH and relative to the corresponding experimental 6-hour EGM2min media, and the mean and SD were determined from the technical triplicates for each sample. Both independent experiments for all isolates were performed for a 6-hour timepoint.

### Binding of Parasite Isolates to HBMEC under Flow Conditions

The binding assay was performed using the Cellix microfluidics system as previously described ^26,29^. Briefly, HBMEC or HDMEC were stimulated overnight with 10 ng/ml TNF, detached, and seeded in attachment factor coated Vena8 biochips (Cellix). When cells formed a confluent monolayer, an IE suspension of 2% parasitaemia and 5% haematocrit in binding buffer (RPMI 1640 with 25 mM HEPES, 11 mM glucose, 2 mM glutamine, pH 7.2) was flowed through for 5 minutes at 37 °C at a shear stress of 0.4 dyne/cm2. After a washing step, bound IE were counted by microscopy in 15 areas throughout the biochip and the mean IE/mm^2^ EC cell surface calculated.

### Statistical Analysis

Statistical analyses were performed on Prism 9 (Version 9.5.1, GraphPad Software). The exact statistical tests, along with the levels of significance, are detailed in the figure legends.

### Data Availability

Data will be made publicly available upon publication.

## Results

### Expression of intermediates involved in maintaining vascular barrier integrity is increased in HBMEC in response to patient-derived *P. falciparum* parasites

To investigate endothelial responses to the patient-derived parasite isolates, we compared the relative gene expression of TNF-activated HBMEC after six-hour exposure to either RBC or IE derived from patients with clinically defined UM or CM. Additional media-only samples were collected to be used as normalization controls in the relative gene expression calculations.

The relative gene expression of 49 target genes was analysed by grouping HBMEC samples by clinical type of the patient isolates, either HBMEC-UM or HBMEC-CM, and comparing them to the corresponding HBMEC-RBC control. Gene expression was calculated for HBMEC-UM, HBMEC-CM, and corresponding HBMEC-RBC by determining the mean FC±SEMs for each group relative to the corresponding 6-hour EGM2min media controls in each of the two independent experiments. Welch’s t-tests were performed to compare the relative gene expression of the HBMEC-IEs to their corresponding HBMEC-RBC controls, and the mean FC±SEMs are reported in Supplementary Table 1.

Differential gene expression was observed in HBMEC in the presence of IE compared to the corresponding RBC control for a number of genes, and several were selected for further study due to their relatively increased expression (FC > 1.7), as well as the biological processes they are involved in. These genes encode: B-cell lymphoma 2-related protein A1 (*BCL2A1*), cytochrome P450 family 1 subfamily A member 1(*CYP1A1*), Krüppel-like factor 2 (*KLF2*), Krüppel-like factor 4 (*KLF4*), prostacyclin synthase (*PTGIS*), and prostaglandin-endoperoxide synthase 2 (*PTGS2*).

A significant increase in expression of *BL2A1*, *CYP1A1*, *KLF2*, *KLF4*, *PTGIS*, and *PTGS2* was observed for HBMEC after co-culture with both clinical isolate types as compared to the RBC controls (Figure 1, Supplementary Table 1). The mean HBMEC gene expression was compared in two ways: grouping by clinical syndrome compared to the corresponding RBC and grouping by combining all IE samples compared to the corresponding RBC. The results for each gene are shown in Figure 1, with a sub-figure showing the combined data for HBMEC-RBC and HBMEC-IE co-culture, with paired data points from the same experiment joined by a line. This allows visual verification of increased gene expression within each experiment, which corresponds to the increase in gene expression observed when samples are grouped by clinical type.

**Figure 1:**
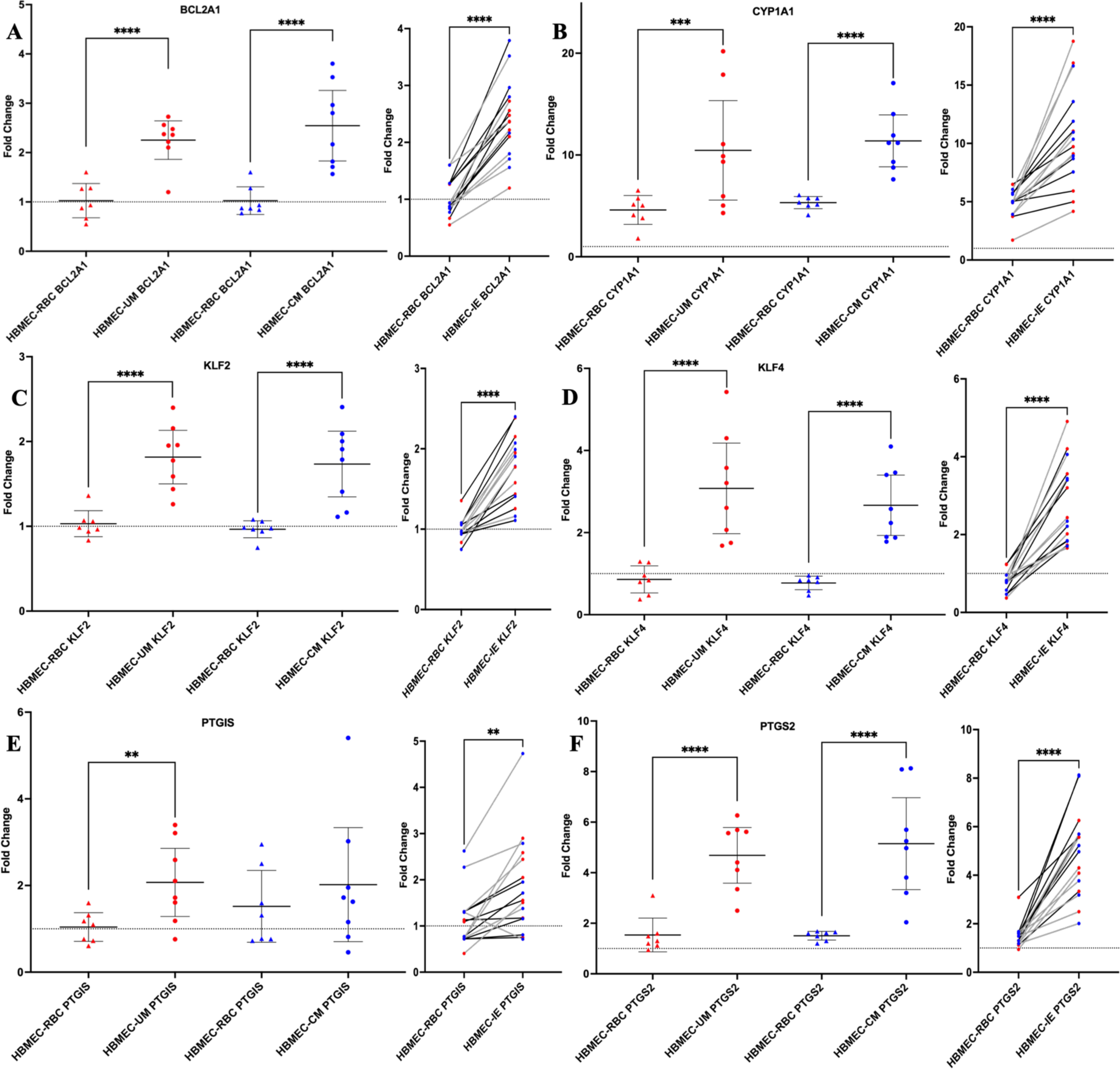
IE of both UM and CM clinical isolates affect HBMEC transcriptional responses as compared to RBC. Differential gene expression of HBMEC after 6-hour co-culture with patient-derived parasites compared to their corresponding RBCs for *BCL2A1*(A), *CYP1A1*(B), *KLF2*(C), *KLF4*(D), *PTGIS*(E), and *PTGS2*(F), calculated as relative to the corresponding 6-hour HBMEC-media control. Mean HBMEC gene expression (Fold Change) was compared in two ways for the selected genes: (left) Welch’s t-test comparing HBMEC-RBC and HBMEC-IE for UM or CM isolates and (right) paired t-test comparing all RBC controls and the combined HBMEC-IE of all isolates. The Welch’s t-test results shown in the graphs are from Supplementary Figure1. Each point on the plot represents one sample FC, calculated as a mean of the technical triplicates, and the mean and 95% confidence intervals are indicated for each group. For the paired analysis, the mean FC was calculated from technical triplicates for each IE and corresponding RBC sample and are coloured red for UM isolates and blue for CM isolates. A paired t-test was performed comparing the mean FC of HBMEC-IE with the corresponding HBMEC-RBC. Lines between the groups indicate that the RBC sample on the left was the control for the corresponding isolate sample on the right. Black lines represent samples from experiment 1, and grey lines represent samples from experiment 2. Two-tailed tests were performed for both analyses, and the dotted line in the plots marks the baseline of FC=1. Significance is depicted by the P-value: *,0.01-0.05; **, 0.001-0.01; ***, 0.0001-0.001; ****, <0.0001.

To validate the observed changes in gene expression, we assessed whether the Fluidigm panel would detect gene upregulation after incubating HBMEC overnight with known stimuli, IL-1beta and TNF, and comparing the relative gene expression with a media control (Supplementary Table 2). Indeed, incubation with these stimuli resulted in upregulation of HBMEC expression in the majority of tested genes, with large fold changes observed in both treatment groups for the following (IL-1beta and TNF, respectively): *CXCL3* (1305 and 862), *ICAM1* (16 and 25), *PTGS2* (30 and 16), *SELE* (1305 and 862), and *VCAM1* (548 and 873).

Previous studies have demonstrated that TNF differentially regulates transcriptional effects on brain endothelial cells ^23,30^, such as upregulated expression of *BCL2A1* in vascular endothelial cells ^31^, 0-hour data was collected to assess if the observed HBMEC FC expression was due to either activation with TNF or the effect of IE. To demonstrate the impact of TNF withdrawal, the relative gene expression of activated HBMEC after 2-hours (0-hour timepoint) and 8-hours (6-hour timepoint) incubation with media were compared for representative samples (Supplementary Figure 1). There was significant reduction of relative FC levels of *KLF4*, *PTGS2*, *ICAM-1*, *VCAM-1*, and an observed reduction in *BCL2A1* in HBMEC after withdrawal of TNF after 8 hours. These findings show that the differences in relative gene expression observed after co-culture are not artifacts of overnight TNF stimulation of the HBMEC.

### *P. falciparum* parasites derived from uncomplicated or cerebral malaria cases do not induce divergent HBMEC responses for the genes tested

To determine whether the observed effect on HBMEC responses to the patient-derived IE was specific to the clinical syndrome, the effect of RBCs on HBMEC was normalized. To take into account variation in the RBCs for each experiment, gene expression FCs (2^−ΔΔCt^) were calculated relative to the corresponding 6-hour HBMEC-RBC sample, which was used as the normalization control (ΔΔCt). Mean gene FCs were calculated for each clinical isolate type (HBMEC-UM and HBMEC-CM) using the triplicates of all the experimental samples. Unpaired t-tests were performed to compare the mean relative gene expression of HBMEC-UM to HBMEC-CM (Figure 2). Apart from *PTGIS*, there was no differential effect on HBMEC transcriptional responses when comparing UM and CM-derived isolates.

**Figure 2:**
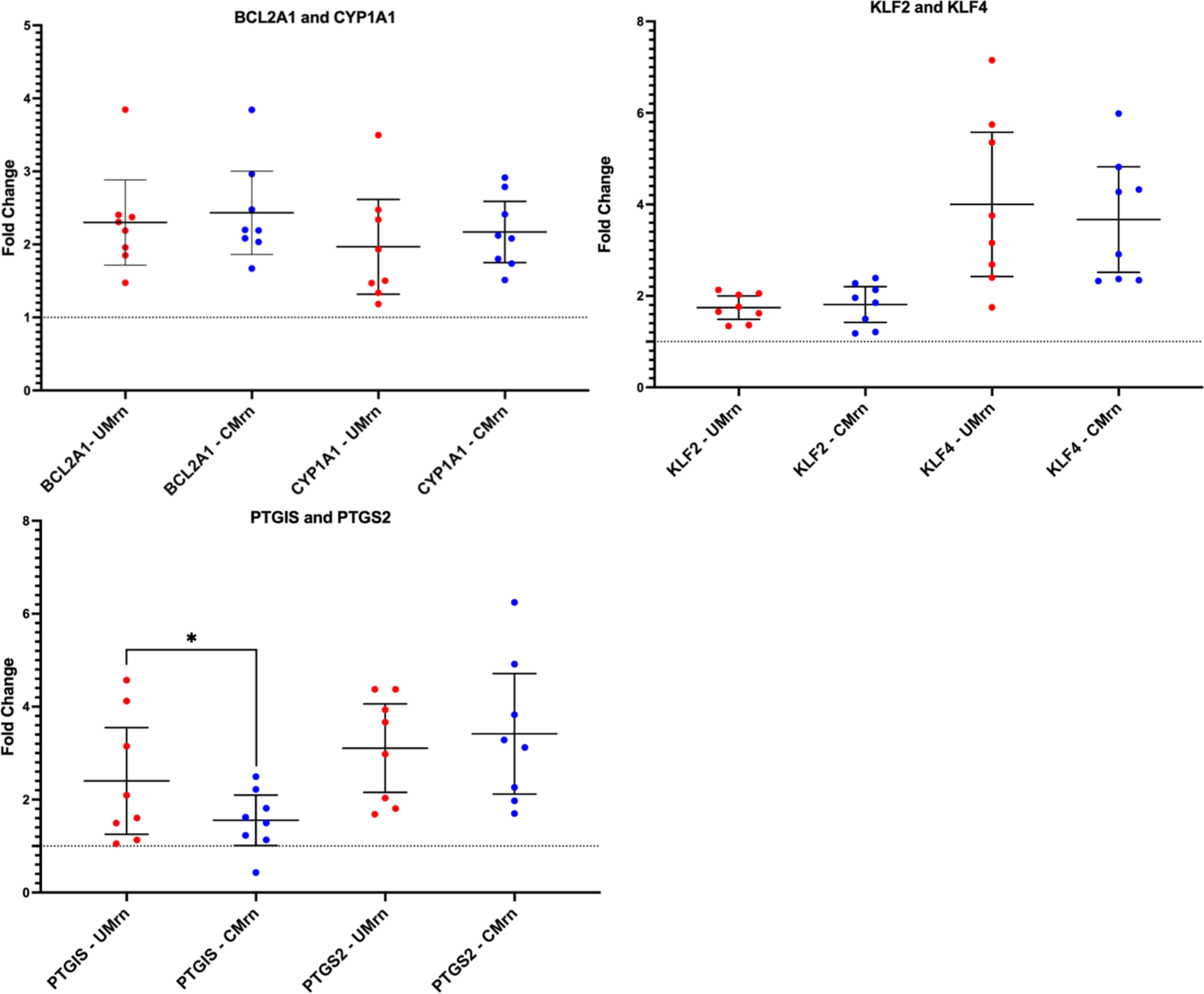
When normalized to HBMEC-RBC controls, isolates within clinical categories do not induce differential responses in HBMEC. To determine the effect of the parasites, HBMEC relative gene expression when exposed to UM and CM patient-derived isolates was calculated relative to HBMEC-RBC expression, using GAPDH as the endogenous control and the 6-hour HBMEC-RBC as the normalization sample reference. Each point represents one experimental sample (mean of triplicates), and the mean FC and 95% confidence intervals for each gene was calculated using both biological replicates (n= 8). RBC normalized HBMEC-UM is represented as “UMrn”, and RBC-normalized HBMEC-CM is represented as “CMrn”. Significance between the two clinical categories was determined by two-tailed unpaired T-tests with P-value summary defined =*,0.01-0.05. The dotted line marks the baseline of FC=1 (red = UM isolates; blue = CM isolates).

The isolates chosen for this study were from a panel of patient-derived *P. falciparum* parasites, isolated from paediatric uncomplicated and cerebral malaria cases in Malawi as described in Storm et al ^26^. During extended periods of culture, and when establishing *in vitro P. falciparum* culture lines from patient parasite isolates, the *var* gene transcription profiles may change which could affect phenotypic changes, such as binding to HBMEC ^32,33^. To establish the isolates in culture and expand the volume to sufficient levels, the experiments were carried out after culturing the IE for 21-60 days. To determine if their adherence to HBMEC had changed due to *var* gene switching over time ^32,33^, binding assays under flow conditions were performed as close to the length of culture time (days in culture, DIC) for each co-culture experiment (Figure 3). Compared to their initial binding phenotype, all isolates showed altered binding to HBMEC. UM1, UM3, UM4, CM2 and CM3 increased their binding after various days in culture, with some variation between the two sets of DIC, and UM2 and CM4 decreased their binding. The SBP1-KO strain does not bind to HBMECs.

**Figure 3:**
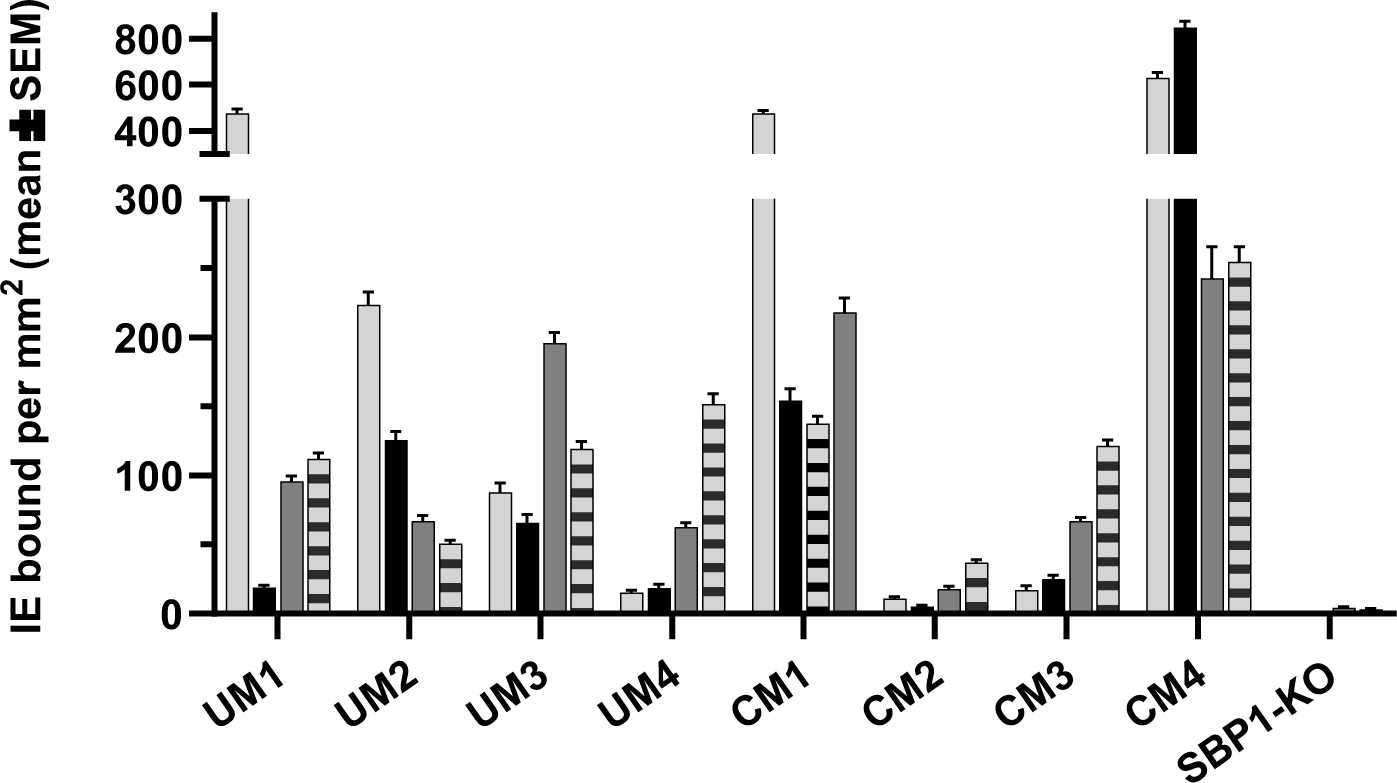
Binding of patient-derived isolates to HBMEC under flow conditions. Binding of IE to TNF-stimulated HBMEC or HDMEC was determined using microfluidics. Shown is the mean ± SEM IE/mm^2^ EC cell surface of 15 fields counted by microscopy. The level of binding for the UM and CM-derived isolates shortly after isolation from the patients is depicted in light grey (HDMEC) and in black (HBMEC). After certain days in culture (DIC) binding of HBMEC is depicted in dark grey (experiment 2) and light grey with horizontal bars (experiment 1). UM1: 30 and 64 DIC, UM2: 32 and 50 DIC, UM3: 40 and 44 DIC, UM4: 25 and 38 DIC, CM1: 31 and 36 DIC, CM2: 32 and 46 DIC, CM3: 50 and 66 DIC, CM4: 42 and 46 DIC. SBP1-KO does not bind to HBMEC.

Using data from both experiments, the binding capacities of the isolates were then correlated with the mean gene expression (FC calculated using the 6-hour RBC as the normalization control) of HBMEC following co-culture with IE using Spearman’s correlation. These calculations were performed for the entire gene panel by grouping the data per experiment, and representative plots for *KLF4*, *PTGIS*, and *PTGS2* are presented in Supplementary Figures 2A and 2B for both experiments. After co-culture, there was no observed significant correlation between the binding avidity of the parasites and the HBMEC gene expression.

### Observed differential effects of *P. falciparum* parasites on HBMEC responses is not solely due to binding

To further investigate whether the observed differential effects on HBMEC transcriptional responses were due to the direct interaction between the IE and the surface of HBMEC, two co-culture experiments were performed using the non-binding parasite strain, skeleton-binding protein knockout parasites (SBP1-KO). These parasites are unable to traffic *Pf*EMP1 to the infected erythrocyte surface, and therefore, they are unable to bind to endothelium ^27,34^, verified by using flow binding assays to HBMEC (Figure 3). The mean relative gene expression was calculated for the technical and biological replicates. Welch’s t-tests were performed to compare the relative FC of the HBMEC-SBP1KO parasites and the HBMEC-RBC controls (Supplementary Table 3). The relative gene expression of the HBMEC incubated with the binding patient-derived parasites (HBMEC-binding IE), and non-binding SBP1-KO parasites, as compared to the HBMEC-RBC controls, are shown in Figure 4 for *BCL2A1*, *CYP1A1*, *KLF2*, *KLF4*, *PTGIS*, and *PTGS2*.

**Figure 4:**
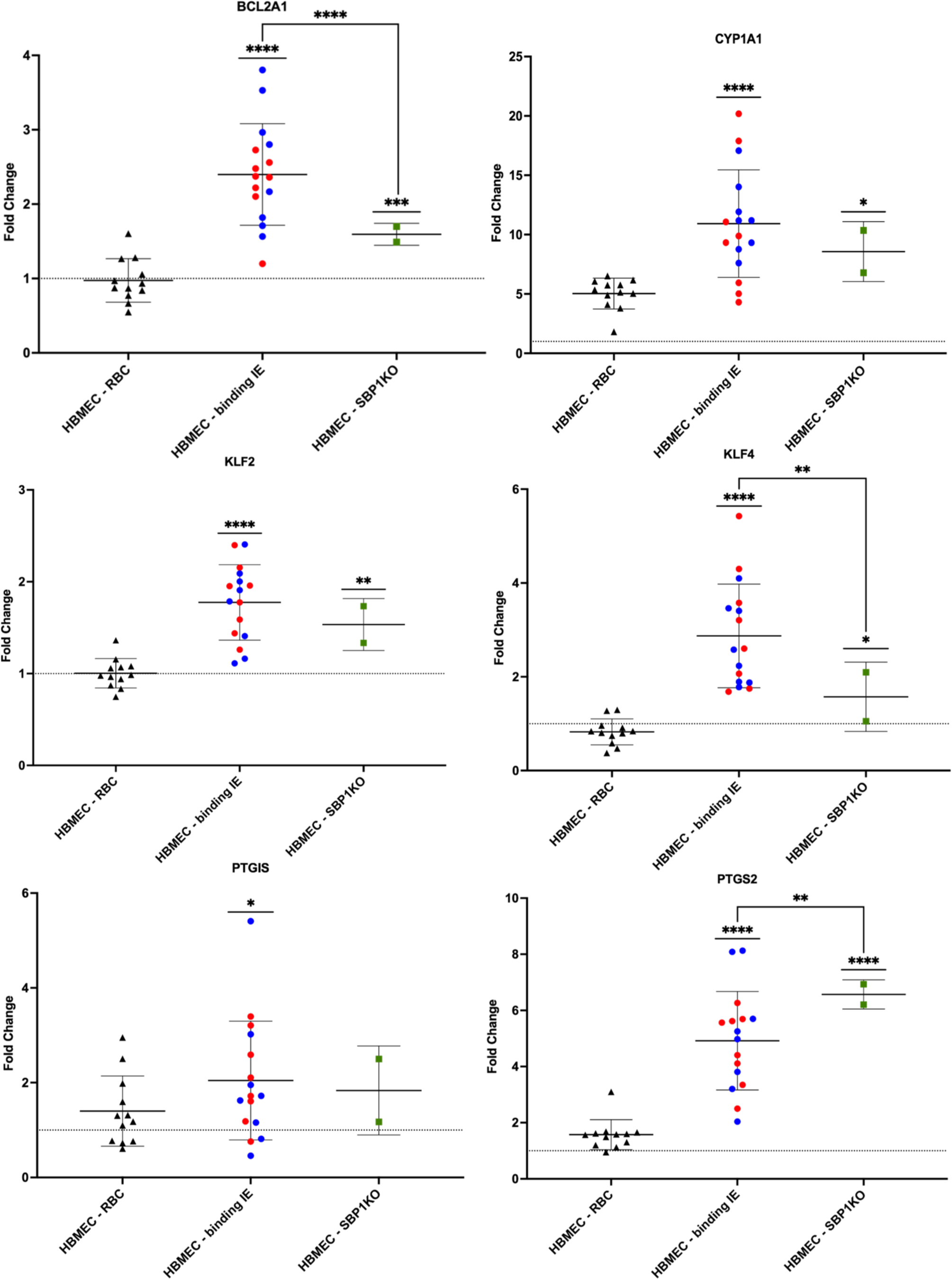
Observed differential effects on HBMEC transcriptional response is not due to binding to HBMEC. Relative gene expression of HBMEC exposed to patient-derived parasites that bind to HBMEC and the non-binding SBP1-KO parasites for 6 hours compared to corresponding RBCs for select genes. HBMEC-RBC samples for both binding and non-binding IE were combined for plotting. Each point on the plot represents one sample FC, calculated as a mean of the technical triplicates, and the mean and 95% confidence intervals are indicated for each group. FC was calculated using GAPDH as an endogenous control and relative to the 6-hour HBMEC-media normalization control (baseline is FC =1). Welch’s t-tests were used to compare HBMEC-RBC and HBMEC-IE for each of the parasite groups, and the mean FC ± SEM for each gene was calculated using the two biological replicates for 4 UM and 4 CM isolates and their corresponding RBC (HBMEC-RBC (binding) n = 10, HBMEC-binding IE n =16, HBMEC-RBC(SBP1KO) and HBMEC-SBP1KO (n =2). The results of the Welch’s t-tests comparing the HBMEC-IE groups with corresponding HBMEC-RBC are summarized in Supplementary Table 3. Sample grouping is represented by symbol, with triangles for HBMEC-RBC, circles for HBMEC-binding IE, and squares for HBMEC-SBP1KO. For binding IE, the red data points represent UM samples, while the blue data points represent CM samples. Samples from experiments with the non-binding parasites are in green. Significant relationships between HBMEC-IE groups and the corresponding HBMEC-RBC are indicated by straight bars over the IE groups (p<0.05), and the P-value summaries are from the calculations performed in Supplementary Table 3. For *BCL2A1*, *KLF4*, and *PTGS2*, the results of Welch’s t-tests comparing HBMEC-binding and HBMEC-SBP1KO are indicated by the bent lines connecting between these groups (p<0.05), and the P-value summaries are from the calculations performed in Supplementary Table 4. In all plots, the dotted line marks the baseline of FC=1. P-value summary defined= *,0.01-0.05; **, 0.001-0.01; ***, 0.0001-0.001; ****, <0.0001).

Even though similar effects were observed from the SBP1-KO parasites on HBMEC transcriptional response when compared to the patient-derived isolates for these genes, the magnitude of the effect is not always the same between HBMEC-SBP1-KO and the HBMEC-binding IE. The relative FC of the HBMEC-binding parasites and the HBMEC-SBP1KO parasites were compared (Supplementary Table 4), and out of the genes presented in Figure 4, there were significant differences between the two groups for *BCL2A1*, *KLF4*, and *PTGS2*. There is large variation in responses within the binding-IE group, likely due to the expression of different *Pf*EMP1variants, suggesting that more than one process is occurring, one of which is adhesion independent. The observed differential effect on HBMEC transcriptional responses is related to parasite-infected RBC, but not exclusively binding via *Pf*EMP1.

As shown in Figure 3, differences in cytoadhesion of the patient-derived isolates to HBMEC were measured after culturing. To determine PfEMP1 domains expressed, *var* genotyping by qPCR was used and the data compared to the qPCR data from when the parasites were isolated from the patients (Supplementary Table 5) ^26^. Primer sets detecting multiple group A and A/B var domains were used, with the CIDRα1 domains predicting PfEMP1 binding to EPCR and the DBLβ domains predicting binding to ICAM-1, both these receptors are expressed on HBMEC. A transcription unit of at least 32 denotes a transcription level equal to the endogenous genes seryl-tRNA synthetase and aldolase, used to normalise the data. High transcript levels of the initial DBLα domains remained for UM4 and CM1 after 26 and 29 DIC, respectively, while the levels decreased substantially for CM2 (43 DIC) and CM4 (33 DIC), indicative for *var* gene switching. CM1 seemed to maintain similar transcripts levels for most of the var domains.

### Determination of prostaglandin endoperoxide synthase 2 and prostacyclin production in HBMEC-IE co-cultures

HBMEC-IE co-culture activates the prostacyclin pathway as shown for *PTGIS* and *PTGS2* in the Fluidigm results (Figure 1). Whether that results in increased concentrations of its end-product prostacyclin or the intermediate PTGS2 was determined by ELISA. Prostacyclin is secreted but has a short half-life, therefore its hydrolysis product, 6-keto prostaglandin F1α (6-keto PGF1α) was measured in co-culture medium of HBMEC-IE and compared with their respective HBMEC-RBC controls (Supplementary Table 6 and Supplementary Figure 3). Overall, there is some variability between the experiments, but no significant differences were detected between the co-cultures. However, the positive controls, HBMEC activated with TNF or thrombin, did increase 6-keto PGF1α concentrations. PTGS2 was detected in HBMEC cell lysate after co-culture and calculated as ng PTGS2 per mg total lysate protein (Supplementary Table 6). Overall, there was a small decrease in PTGS2 generation after HBMEC-IE co-culture, but only significant for CM1 (experiment 9) and CM4 (experiment 6).

### Production of cytokines by HBMEC-IE co-cultures

To determine if the production of cytokines or chemokines by HBMEC was altered following co-culture, a multiplex panel of 41 secreted cytokines and chemokines was used. The multiplex was performed at an early stage of the study and only included 6-hour co-culture medium of five isolates and two lab strains IT4var14 and IT4var37 in one experiment. These lab strains were included because they were used in previous co-culture experiments ^25^. Concentrations higher than 5 pg/ml were detected for 13 cytokines and only a few of these were significantly increased after co-culture with IE, compared to co-culture with their respective RBC control (Supplementary Table 7). CM4, the isolate with the highest binding to HBMEC, significantly increased production of IL-6, IL-8 and IP-10, GRO, G-CSF and GM-CSF. Although at low levels, IFNγ was increased two to ten-fold by co-culture with the patient isolates, but not significantly by the lab strains IT4var14 and IT4var37. A positive control of TNF-activated HBMEC was included to produce high enough levels of cytokines to be measured, and for 10 analytes these were at least 10-fold higher than medium only. Notably, TNF was still detectable at ∼11 ng/ml in the culture medium after 16 hours.

### Effect of IE co-culture on HBMEC barrier integrity

Although the cytoadherence phenotype of the isolates to HBMEC altered during culture, there are still detectable differences, especially between CM2 and CM4 with low and high binding, respectively. Whether differential binding capacity of the patient isolates could affect HBMEC barrier function was determined by trans endothelial electrical resistance (TEER). Barrier function increased slightly by adding RBC, irrespective if they were infected or not (Supplementary Figure 4B). Thrombin induces a rapid decrease in barrier function, which recovers within 2 hours. The presence of RBC and IE partly protected against this effect by reducing the maximum decrease (Supplementary Figures 4C and 4E) and the rate of recovery, determined by the area under the curve (AUC) (Supplementary Figures 4D and 4F). RBC and IE all significantly reduced thrombin-induced decrease in barrier function compared to medium, but IE was not significantly different to RBC. The recovery rate was more rapid with a significantly reduced AUC for RBC, 5%, 10%, and 30% IE compared to medium with a trend of more rapid recovery with increasing parasitaemia. The AUC for SBP1-KO at the different parasitaemias were similar and not distinct compared to medium, but all significantly different than the AUC of 30% IE, which had the smallest AUC at 13.5. No differences between the individual UM and CM-derived isolates could be detected.

## Discussion

A view of CM pathology has emerged in recent years that proposes dysregulation of coagulation pathways, inflammation, release of parasite factors from mature IE, and sequestration of IE in the vasculature all contribute to endothelial dysfunction and breakdown of the BBB ^8,9,35^. Cytoadhesion of IE to endothelial cells is one of the events in a multi-step process leading to localized accumulation of both IE and RBC, as well as rupture and release of parasite and intercellular erythrocyte components and soluble factors, resulting in activation of the endothelium and immune responses, compromising BBB integrity ^20,36^.

In the present study, HBMEC responses to clinically derived parasites isolates from UM and CM patients were investigated by employing an *in vitro* co-culture model of cytoadhesion. We found that IE from both UM and CM-derived isolates have differential effects on the HBMEC transcriptional response as compared to RBCs for 28 genes (Supplementary Table 1 and Figure 1), but there was no difference when comparing UM with CM patient-derived isolates (Figure 2). The UM and CM-derived isolates were selected for this study based on their diverse binding capacities to HBMEC and HDMEC from a previous study ^26^, and their ability to be established in continuous culture to provide sufficient material for repeat experiments. Adapting *P. falciparum* patient-derived isolates to *in-vitro* culture requires considerable time, and thus *var* gene switching is likely to occur ^32,37^, which was the case for the CM and UM isolates, as detected by qPCR (Supplementary Table 5). This resulted in altered binding profiles to TNF-stimulated HBMEC with less distinction between the isolates compared to that seen at the time of isolation from patients, with exceptions of CM2 and CM4, which still had relatively low and high binding to HBMEC, respectively (Figure 3). Binding to HDMEC was not determined in this work for the cultured-adapted isolates, as the co-cultures were only performed with HBMEC. Despite the variability in binding profiles, we observed important differences in the HBMEC transcriptional response to IE compared to RBC in our *in vitro* model, supporting a role for PfEMP1 in shaping the endothelial response to sequestration.

Comparing the binding capacity of each isolate to the relative gene expression of HBMEC after 6-hours co-culture detected no significant correlation between the parasites’ propensity to bind and the HBMEC gene expression of *PTGIS*, *PTGS2* or *KL4* (Supplementary Figure 2). Furthermore, experiments performed using the SBP1-KO parasite line revealed that there were no differences in the HBMEC responses to binding and non-binding IE for these genes (Supplementary Table 3), yet there was a significant difference in the magnitude of these responses between these two groups for 30 genes (Supplementary Table 4). SBP1-KO parasites do not cytoadhere to HBMEC as they are unable to traffic PfEMP1 to the surface, although they maintain both Maurer’s cleft and knob morphology and the expression of other Maurer’s cleft-associated proteins essential for tracking and IE surface changes, such as KAHRP, MAHRP, REX, PfEMP3, and Pf332 ^27,34^. RIFINs and STEVORs are proteins that are expressed on the surface of IE that are involved in cytoadherence to RBC (rosetting) and immune cells ^38^, and their trafficking and expression is maintained in SBP1-KO parasites ^39^. Possemiers et al investigated sequestration-deficiency in experimental *Plasmodium berghei* malaria-associated acute respiratory distress syndrome, and they observed endothelial activation in SBP1-KO infected mice to the same level as wild type at eight days post infection in the lungs^40^. Overall, SBP1-KO IE maintain an altered surface, including knobs, and can interact with HBMEC differently compared to RBC. Our findings suggest that the interaction between HBMEC and IE via PfEMP1 is not the only driver and that the observed differences in HBMEC relative gene expression are also related to contact with intact parasitised RBC.

We observed upregulation in relative gene expression of HBMEC after incubation with IE in a panel of genes, particularly in genes involved in apoptosis and the regulation of inflammation and endothelial integrity. Overall, the fold change expression levels were relatively low, except for the ones highlighted for further study and discussion. *BCL2A1* is a member of the pro-survival sub-family of BCL2 proteins that include both pro- and anti-apoptotic regulators, and it encodes a protein that reduces the release of pro-apoptotic cytochrome C and blocks caspase activation through inhibition of caspace9 ^41,42^. Expression of *BCL2A1* is upregulated in vascular endothelial cells by the inflammatory cytokines TNF and IL-1beta ^31,43^. *BCL2A1* expression was increased and a more robust *BCL2A1* expression was observed in HBMEC co-cultured with binding IE, as compared to non-binding SBP1-KO (Supplementary Table 3). Perhaps expression of *BLC2A1* early during an infection represents the balance between protection and pathology and demonstrates the temporal kinetics in this complicated system. KLF2 and KLF4 are transcription factors that have been shown to directly regulate endothelial function and integrity in vascular endothelium and to protect against endothelial dysfunction, both *in vitro* and in murine models, and their expression is modulated by both changes in laminar sheer flow or stress and pro-inflammatory stimuli ^44–49^. Overexpression of *KLF2* and *KLF4* affect coagulation and inflammatory pathways by inducing expression and activity of endothelial nitric oxide synthase and thrombomodulin ^44–46,50,51^, while simultaneously inhibiting expression of adhesion molecules E-selectin, ICAM-1, and VCAM-1 ^45^. We observed significant upregulation in relative gene expression of both *KLF2* and *KLF4* in HBMEC incubated with IE. CYP1A1, PTGIS, and PTGS2 are involved in lipid metabolism, including the biosynthesis of prostacyclin, which has roles as an inflammatory mediator and a potent vasodilator ^52,53^. CYP1A1 is a member of the cytochrome P450 enzyme family that metabolizes a range of substrates, including arachidonic acid ^54^. Both PTGIS and PTGS2 are enzymes involved in the biosynthesis of prostacyclin, a bioactive lipid that is produced by vascular endothelial cells and that plays a role in regulation of endothelial inflammation and apoptosis ^52^.

A recent study demonstrated differential gene expression of *CYP1A1*, *PTGIS*, and *PTGS2* between HBMEC incubated with two parasite lines that expressed different *Pf*EMP1s ^25^. As reported in that study, we observed evaluated levels of CYP1A1 induced by both IE and RBC controls. Despite observing transcriptional upregulation of *PTGIS* and *PTGS2*, FC±SEM of 2.05±0.24 and 4.92±0.26 respectively (Supplementary Table 3), this did not seem to translate into detection of PTGS2 protein or 6-keto PGF1α (prostacyclin’s stable hydrolysis product) in HBMEC lysate and co-culture supernatant, respectively (Supplementary Table 6). PTGS2 is intracellularly localised, and expression in cell lysates is usually detected by Western blot. An ELISA was employed to quantify PTGS2 concentrations in multiple samples. Unfortunately, the usability of the ELISA kit was questionable, as the positive control, stimulation with TNF, did not result in increased PTGS2 production, and detection of PTGS2 was not pursued further. 6-keto PGF1α is secreted, and larger amounts were produced upon stimulation with TNF and thrombin, but overall, no significant increase in production was detected for the HBMEC-IE co-cultures compared to their respective HBMEC-RBC co-cultures. Overall, our findings provide additional evidence that IE regulate brain ECs and that these effects are not only due directly to binding.

Previous studies have investigated the effects of IE on endothelial responses *in vitro*. An investigation by Zuniga et al examined how TNF and IE differently affected transcriptional responses and barrier integrity in brain endothelial cells. Their work highlighted that TNF and IE lysates induce distinct transcriptional profiles, with TNF associated with induction of endothelial activation while IE lysate contributed considerably to endothelial barrier disruption, yet overlap in genes associated with inflammation, including induction of PTGS2, were detected ^23^. Howard et al adapted a 3D perfusion human brain microvessel model to evaluate the brain endothelial responses to perfusion with TNF and at different stages of parasite maturation. This study found that treatment of brain microvessels with TNF induced an inflammatory phenotype, while EPCR-binding parasites caused localised barrier damage, and induced unique stress response pathways associated with metal toxicity and oxidative stress ^30^. In concordance with Zuniga et al and Howard et al ^23,30^, we observed that a group of inflammatory associated genes are upregulated in HBMEC by exposure to patient-derived IE and this effect is independent of TNF treatment. This upregulation in HBMEC relative gene expression was detected after 6-hours incubation with trophozoite-stage IE, in contrast to studies that used schizont-stage IE or IE lysates for longer periods ranging from 6, 9, 12, and 24 hours ^23,30^. In contrast to these studies ^23,30,55^, we utilised trophozoite-IE prepared in EGM2min medium for a 6-hour incubation period. This medium maintains HBMEC over the time course, but affects the progress of IE, as they barely develop beyond the trophozoite stage. In our study, we detect HBMEC effects at early stages of sequestration where IE are still in contact with the endothelium, as opposed to effects attributed to parasite material and ruptured IE. An extended co-culture time frame would have allowed us to observe other, perhaps larger, effects on HBMEC transcription. This agrees with the proposed multi-step activation of endothelium in which binding results in a high local concentration of soluble erythrocytic and parasitic factors ^20^.

A multiplex assay was used to detect 41 cytokines in the co-culture supernatant. HBMEC produce low levels of cytokines in their basal state, and this is not much increased after co-culture with IE, with a relatively low FC comparing IE to RBC co-culture (Supplementary Table 7). The multiplex was not performed with all the isolates nor SBP1-KO, and only with one experiment, thus, the results are only an indication of the effects of IE co-culture. Stimulation with TNF, as positive control, did increase the concentrations of multiple cytokines, including IL-6, IL-8, IP-10 and MCP-1, also reported in other studies ^23,55,56^. Indeed, Zuniga et al^23^ noted they observed transcriptionally similar but translationally different levels of cytokines between their treatment groups even when total protein synthesis levels were similar, suggesting a role for post-translational regulation in HBMEC after incubation with IE.

Whether the patient-derived isolates affect HBMEC barrier integrity was determined by TEER, with cell index as measure of integrity. In all experiments, the cell index was increased after adding IE, but to a similar level as RBC, likely to be due to the layer of RBC on top of the HBMEC affecting conductivity (Supplementary Figure 4A and B). None of the isolates decreased the HBMEC barrier function, even after 24 hours (Supplementary Figure 4B). This contrasts with other studies where the addition of IE decreased the barrier integrity ^23,55^ and is likely related to their use of schizont-stage IE or IE lysates, while trophozoite-stage IE were used in our experiments. After 24 hours, parasites seemed to be in a resting stage and were not progressing to schizont stage, therefore not releasing parasite factors, previously identified as the reason for the decrease in barrier function ^23,55,57^.

It has been reported that IE affect the endothelial responses to thrombin ^23,55^, which reduces barrier function rapidly and quickly recovers. The presence of RBC or IE affected the response of HBMEC to thrombin with a reduced maximum response and AUC, compared to medium (Supplementary Figures 4E and F). There were no significant differences between RBC or IE, but the non-binding lab isolates SBP1-KO, at the three different parasitemias, was significantly different to the effect of 30% IE (Supplementary Figure 4D, E, and F), indicating that receptor binding may be required to reduce the effect of thrombin. In contrast, Avril et al showed that schizont IE, but not trophozoite IE, prolonged thrombin-induced barrier disruption, compared to RBC ^55^. However, thrombin was added 2 – 3 hours after adding IE or RBC and not compared to the effect of thrombin in medium. Thrombin decreases endothelial barrier function by cleavage of Protease Activated Receptor 1 (PAR1) and involves the thrombomodulin /activated protein C system in which the balance of thrombin and activated protein C determines PAR1-dependent barrier disruptive or protective action, respectively ^14,58^. The reduced effect of thrombin indicates a shift towards more activated protein C, which was not investigated in this study. However, binding of IE to HBMEC induces receptor shedding and this could change the level of thrombomodulin and endothelial protein C receptor, the receptors for thrombin and protein C, respectively ^57,59^, affecting PAR1 cleavage. The TEER data show that specific binding of IE to HBMEC do affect certain endothelial responses, not captured in the RNA transcription data.

One limitation of this study was the use of a co-culture model that simplified the complex microvascular system down to static interaction between IE and HBMEC. Nonetheless, this model allowed for the study of the HBMEC transcriptional response to patient-derived parasites and to assess if these responses were dependent on IE cytoadhesion or clinical syndrome. The ratio of 110 IE/HBMEC in our experiments is similar to the range employed in other studies ^23,55^, which was calculated in Zuniga et al to be equivalent to approximately two layers of IE on top of the HBMEC. We tested whether the Fluidigm panel would reflect endothelial gene expression changes by exposing HBMEC overnight to IL-1beta and TNF and comparing the gene expression with a media control (Supplementary Table 2). We observed massive upregulation in HBMEC expression of several genes, including *CXCL3*, *ICAM1*, *PTGS2*, *SELE*, and *VCAM1*, after incubation with both stimuli, confirming that our model was able to determine regulation of specific genes. Furthermore, there was significant reduction in relative gene expression after withdrawal of TNF from HBMEC (Supplementary Figure 1). Overall, these findings suggest that the changes in relative gene expression observed in HBMEC exposed to both binding and non-binding parasites are due to the impact of IE.

During early stages of sequestration, we observed effects on endothelium that are both dependent and independent of *Pf*EMP1. These effects were relatively small except for several genes, including *CYP1A1*, *KLF4*, and *PTGS2*. These genes may represent early endothelial protective responses that act through vascular regulation and modulation of thrombomodulin. There was no difference in gene regulation in HBMEC based on the origin of the isolate nor the parasite binding capacities. Overall, interaction of IE and endothelial cells in early stages of sequestration does induce some endothelial responses, including some that are independent of PfEMP1, and this is the first investigation to our knowledge to study how non-*Pf*EMP1 expressing IE affect endothelial transcriptional responses. Understanding the different mechanisms of crosstalk between *P. falciparum* infected erythrocytes and host endothelium may help us to develop interventions that support patients with severe malaria while effective anti-parasite drugs clear the infection.

## Acknowledgements

We thank the members of staff in the NGS facility at the Bioscience Core Labs in KAUST for the training and use of the Fluidigm Biomark HD system. This work was supported by Competitive Research Grant (CRG) to AP and AGC (Grant number: CRG6-3392.02) by the Office of Sponsored Research (OSR) at the King Abdullah University of Science and Technology (KAUST). The funder had no role in study design, data collection, or preparation of the manuscript.

## Author Contributions

Study conceptualisation: AGC; funding acquisition and administration: AGC, AP; study supervision: AGC, AP, JS; study investigation: CA, JS, GC; formal analysis: CA, JS; writing – original draft: CA, JS; writing – review and editing: CA, JS, AGC, AP. CA and JS determined CA would be listed first as they are an early career scientist and JS is already an established researcher.

## Supplementary Data

**Supplementary Table 1:**
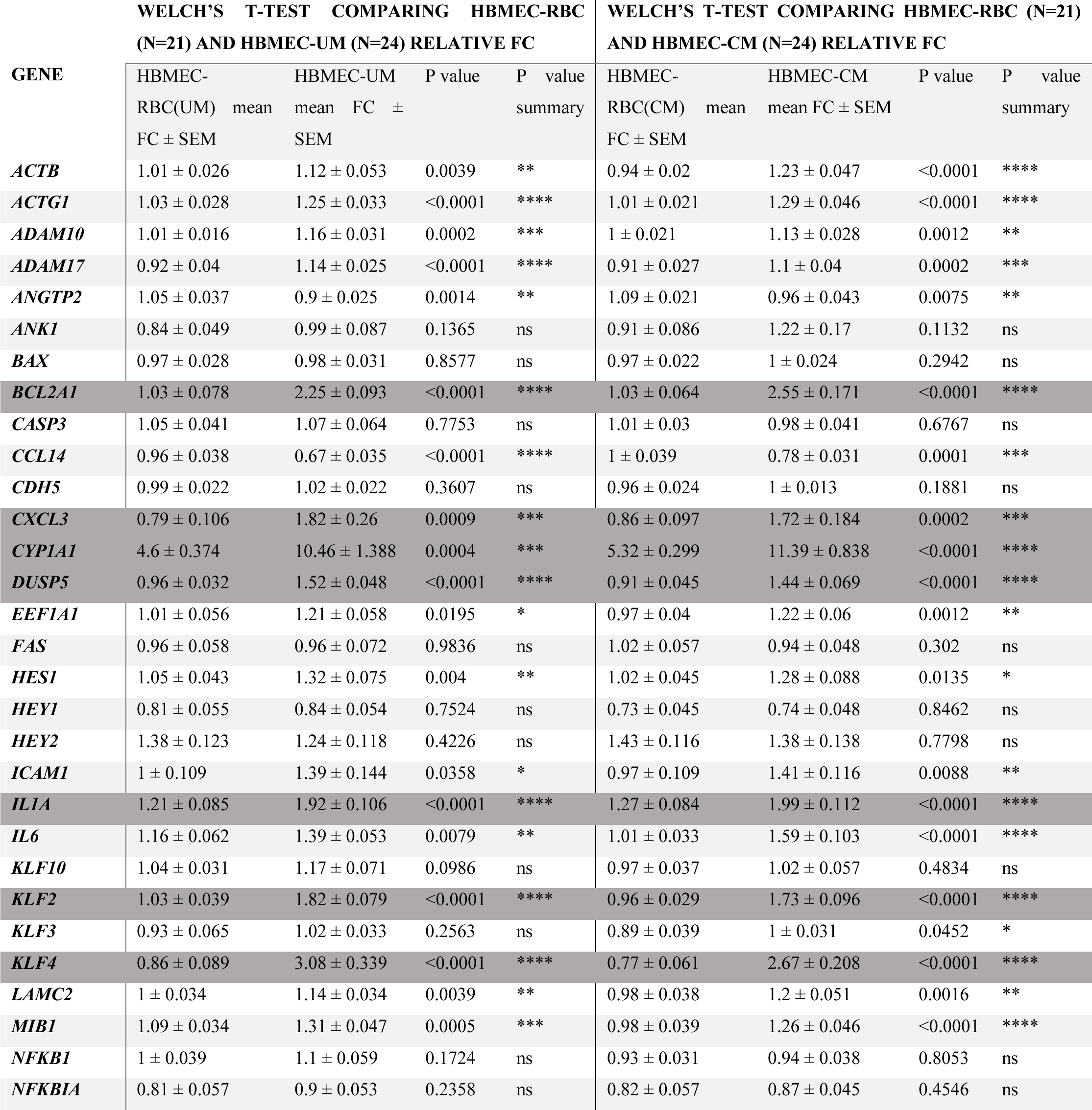
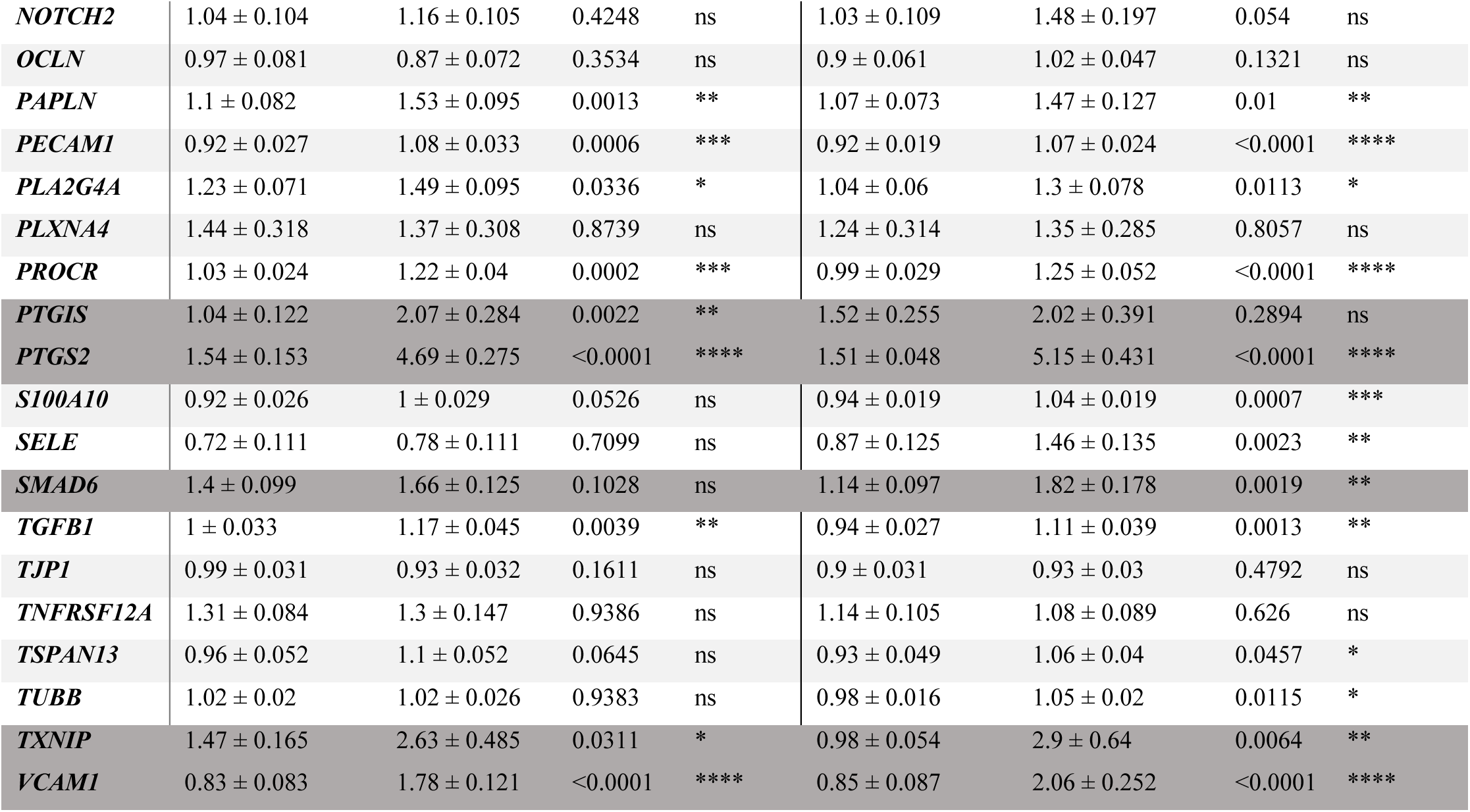
Relative gene expression of HBMEC co-cultured with IE from patient-derived isolates or with the corresponding RBC control for each gene in the panel. Samples were collected after a 6-hour co-culture, and relative gene expression was calculated using the ΔΔCt method using GAPDH as the endogenous control and the corresponding experimental 6-hour HBMEC-media sample as the reference sample (baseline is FC =1). The mean Fold Change (FC) and standard error of the mean (SEM) for each gene were calculated using the technical triplicates of the two biological replicates (UM = uncomplicated malaria patient-derived isolates, CM = cerebral malaria patient-derived isolates, n = sample size). Rows shaded in dark grey have FC>1.5 for one of the IE groups. P-value summary defined= *,0.01-0.05; **, 0.001-0.01; ***, 0.0001-0.001; ****, <0.0001.

**Supplementary Table 2:** Assessment of endothelial responses and Fluidigm panel through relative gene expression of HBMEC after overnight incubation with inflammatory cytokines. HBMEC were incubated overnight with either EGM2min media, 1 ng/mL IL-1beta, or 10 ng/mL TNF. The samples were collected directly after stimulation, and the relative gene expression was determined for several genes.. FCs were calculated using GAPDH as an endogenous control as relative to the HBMEC-overnight media normalization control (baseline is FC=1). The mean FC for each gene was calculated from technical triplicates from one experiment (n=3).

| GENE | HBMEC-OVN MEDIA<br>RELATIVE FC | HBMEC OVN IL-1B<br>RELATIVE FC | HBMEC-OVN TNF<br>RELATIVE FC |
| --- | --- | --- | --- |
| *BCLA2A1* | 1 | 4.7 | 6.0 |
| *CXCL3* | 1 | 1305 | 862 |
| *CYP1A1* | 1 | 1.1 | 0.6 |
| *ICAM1* | 1 | 16 | 25 |
| *IL1A* | 1 | 40 | 30 |
| *IL1B* | 1 | 60 | 5.4 |
| *KLF2* | 1 | 1.1 | 1.0 |
| *KLF4* | 1 | 3.1 | 2.9 |
| *PTGIS* | 1 | 0.8 | 1.2 |
| *PTGS2* | 1 | 30 | 16 |
| *SELE* | 1 | 72 | 52 |
| *VCAM1* | 1 | 548 | 873 |

**Supplementary Figure 1:**
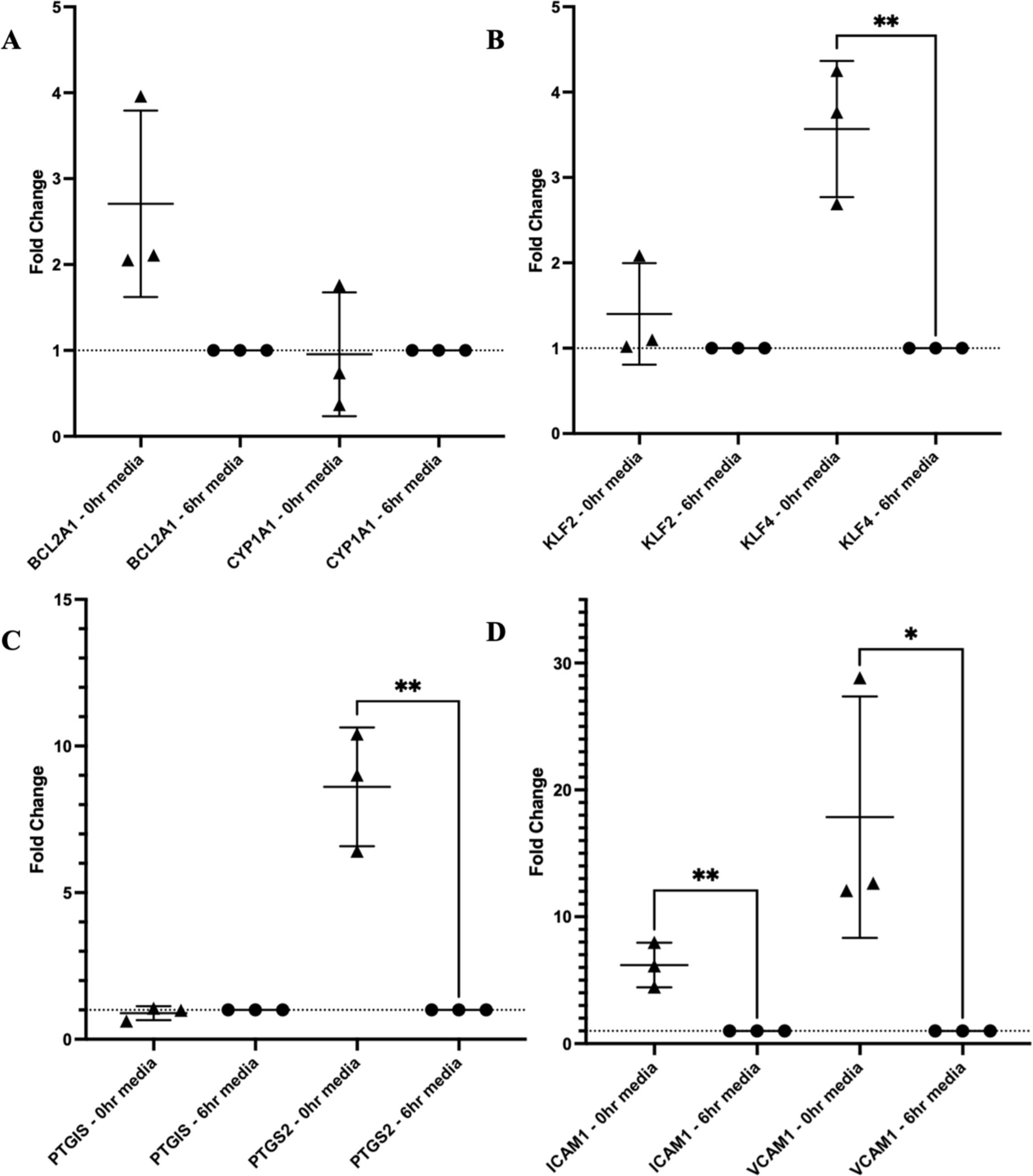
Representative relative gene expression of HBMEC after withdrawal of TNF stimulation. In the co-culture model, HBMEC are stimulated overnight with 10 ng/mL of TNF to stimulate the cells. The next morning, the supernatant containing TNF was removed and replaced with EGM2min media for two hours before co-culture experiments commenced. HBMEC-media samples were collected at 0hr and 6hour culture time points, corresponding to 2- and 8-hours incubation with EGM2min media and without TNF, respectively. Relative gene expression was determined for (A) *BCL2A1* and *CYP1A1*, (B) *KLF2* and *KLF4*, (C) *PTGIS* and *PTGS2*, and (D) *ICAM-1* and *VCAM1* for HBMEC-media samples for three independent, representative experiments (n=3). FCs were calculated using GAPDH as an endogenous control as relative to the HBMEC-6hour media normalization control (baseline is FC=1). Each plot point represents a mean of the technical triplicates, and the mean ± SD of the 3 experiments is shown with triangle points representing HBMEC-0hr media samples and circular points representing HBMEC-6hr media samples.

**Supplementary Figure 2:**
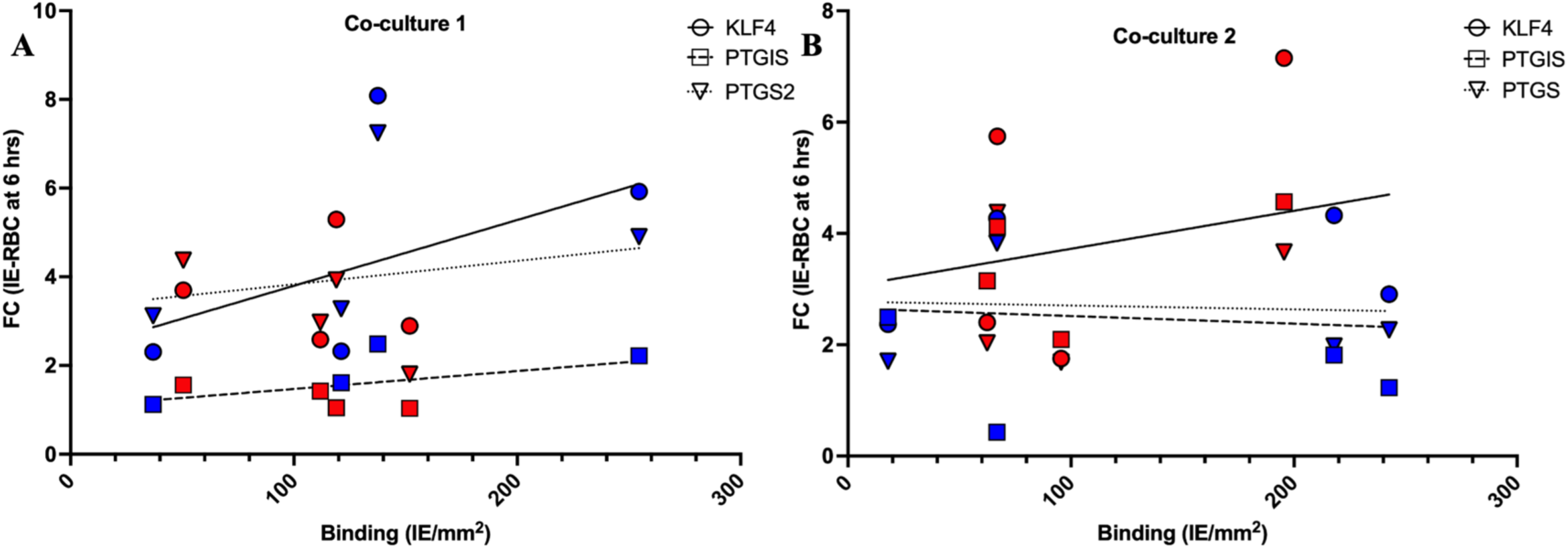
Binding capacities of patient-derived parasite isolates do not correlate with HBMEC gene expression. The binding capacity of each isolate (IE/ mm^2^) was compared to the 6-hour HBMEC mean FC relative gene expression for select genes (red = UM isolates; blue = CM isolates) after the first (A) and second (B) experiments using a linear regression. The linear regression for KLF4, PTGIS, and PTGS2 are indicated on each graph by straight, dashed, and dotted lines, respectively, and the R^2^ values are reported as follows: Co-culture 1) KLF4 = 0.22, PTGIS= 0.25, PTGS2= 0.05; Co-culture 2) KLF4 = 0.10, PTGIS= 0.01, PTGS2= 0.003. Relative gene expression was calculated for HBMEC-IE using the ΔΔCt method with GAPDH as the endogenous control and the corresponding 6-hour HBMEC-RBC as the reference sample (baseline is FC=1). Each point represents the mean FC for each HBMEC-IE sample (n=8), calculated from the technical triplicates. Pearson’s correlation was performed to compare parasite binding capacity to the differential gene expression for each isolate for both experiments, and the correlation coefficient (r) values are reported as follows: Co-culture 1) KLF4 = 0.47, PTGIS= 0.50, PTGS2= 0.22; Co-culture 2) KLF4 = 0.31, PTGIS= −0.08, PTGS2= −0.05.

**Supplementary Table 3:**
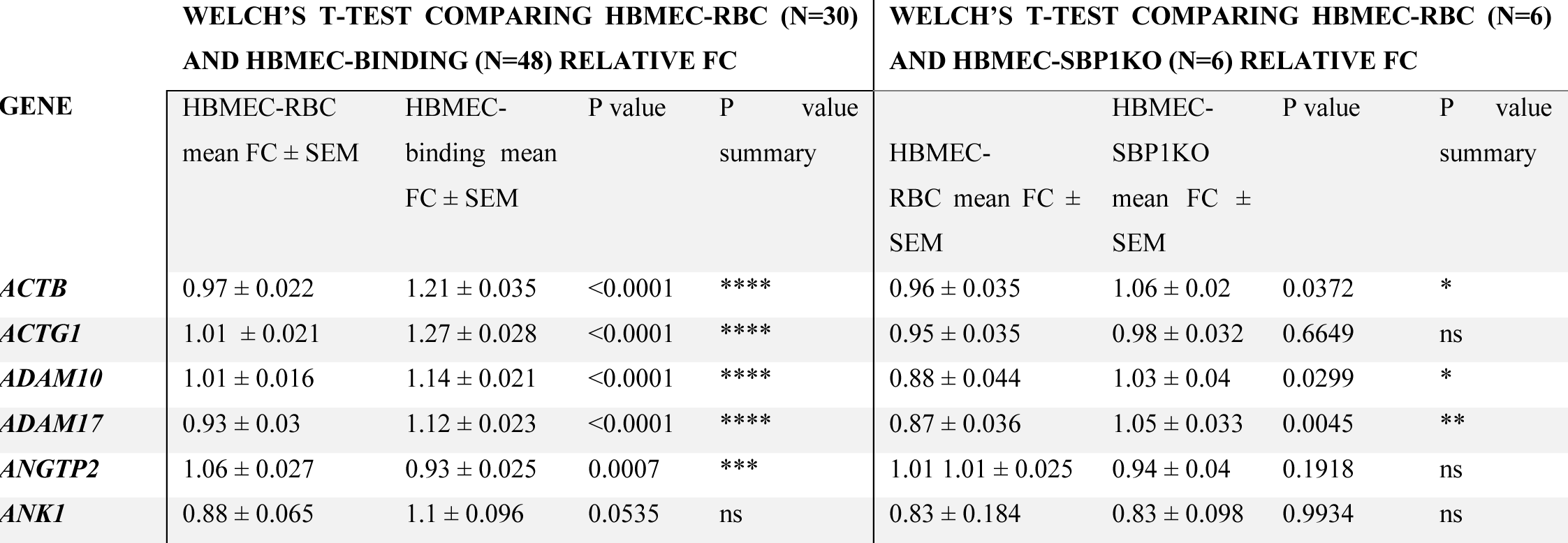
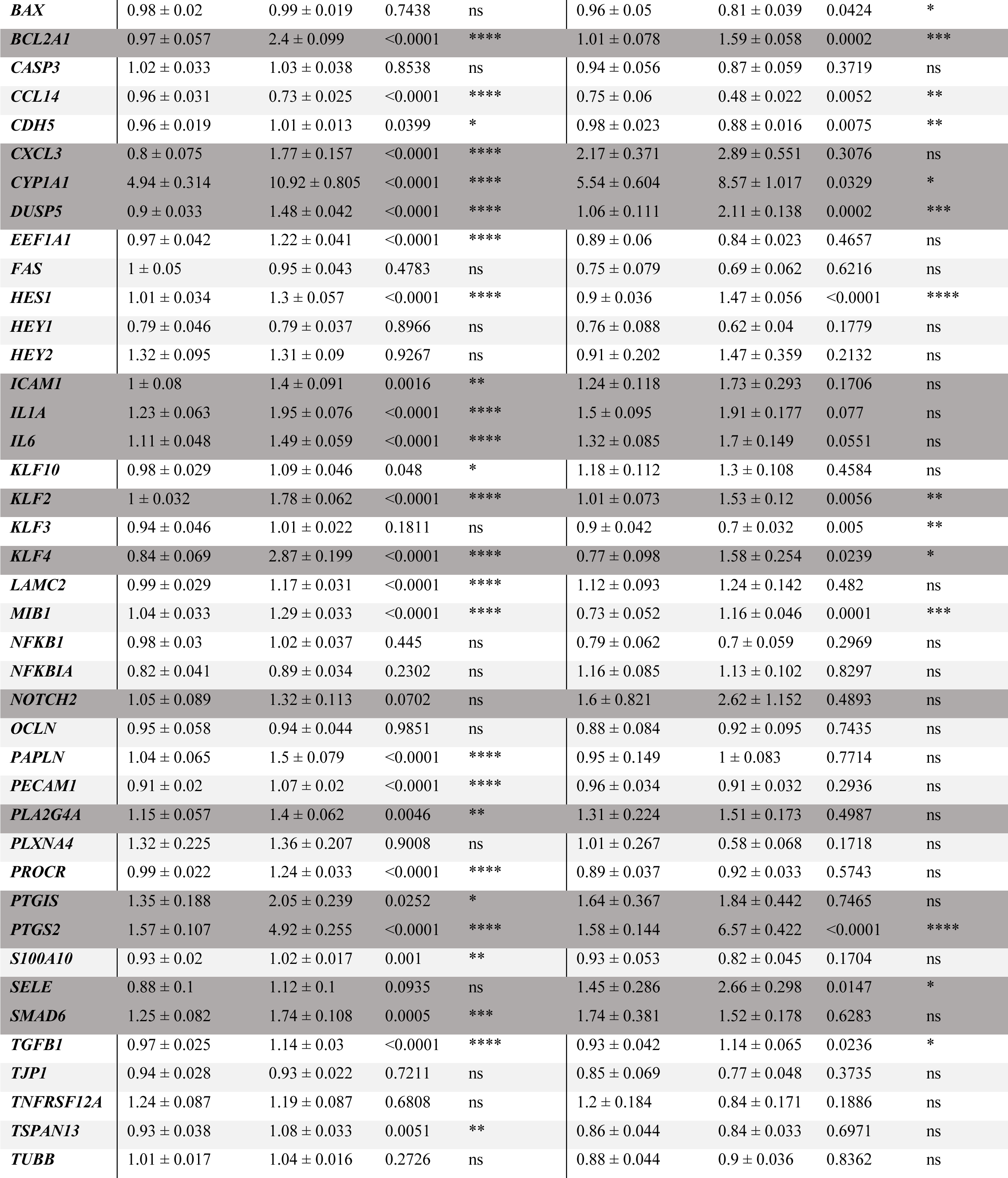
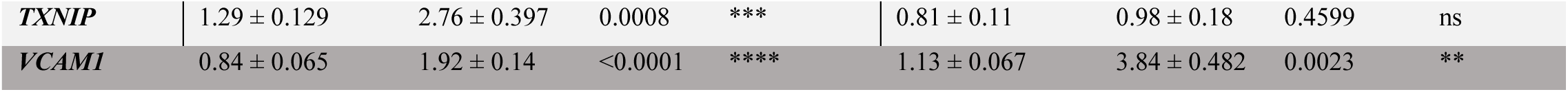
Relative gene expression of HBMEC co-cultured with either binding IE or non-binding IE compared to the corresponding HBMEC-RBC control for each gene in the panel. All samples were from a 6-hour time point. Data from UM and CM patient-derived parasite isolates were combined to form a group of binding parasites, while two experiments with non-binding SBP1KO parasites were performed. Relative gene expression was calculated using the ΔΔCt method using GAPDH as the endogenous control and the corresponding experimental 6-hour HBMEC-media sample as the reference sample (baseline is FC=1). The mean Fold Change (FC) and standard error of the mean (SEM) for each gene were calculated using all experimental samples and technical replicates (SBP1KO= SBP1-knockout parasites, n = sample size). Rows shaded in dark grey have FC>1.5 for one of the HBMEC groups. P-value summary defined= *,0.01-0.05; **, 0.001-0.01; ***, 0.0001-0.001; ****, <0.0001.

**Supplementary Table 4:**
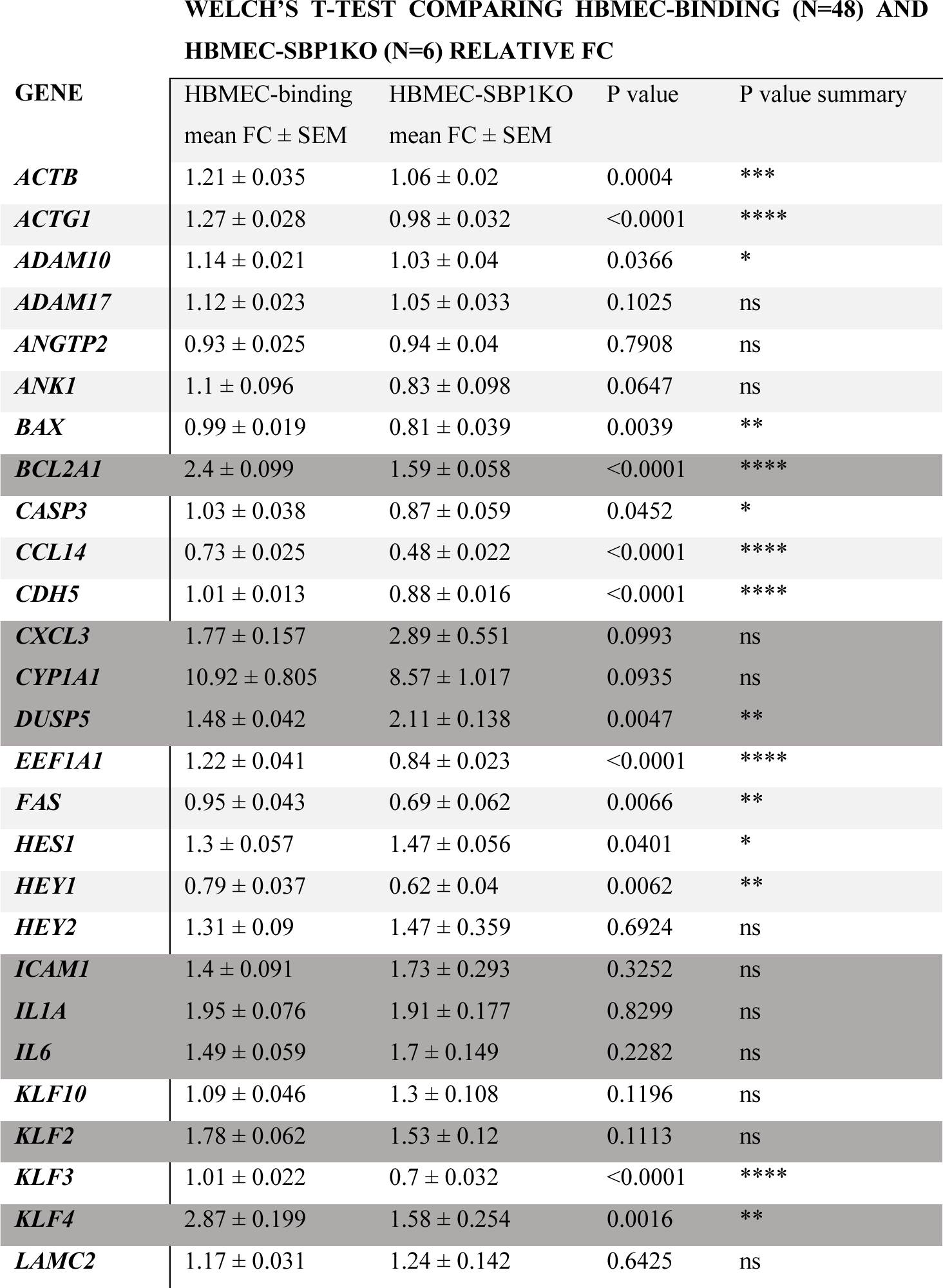
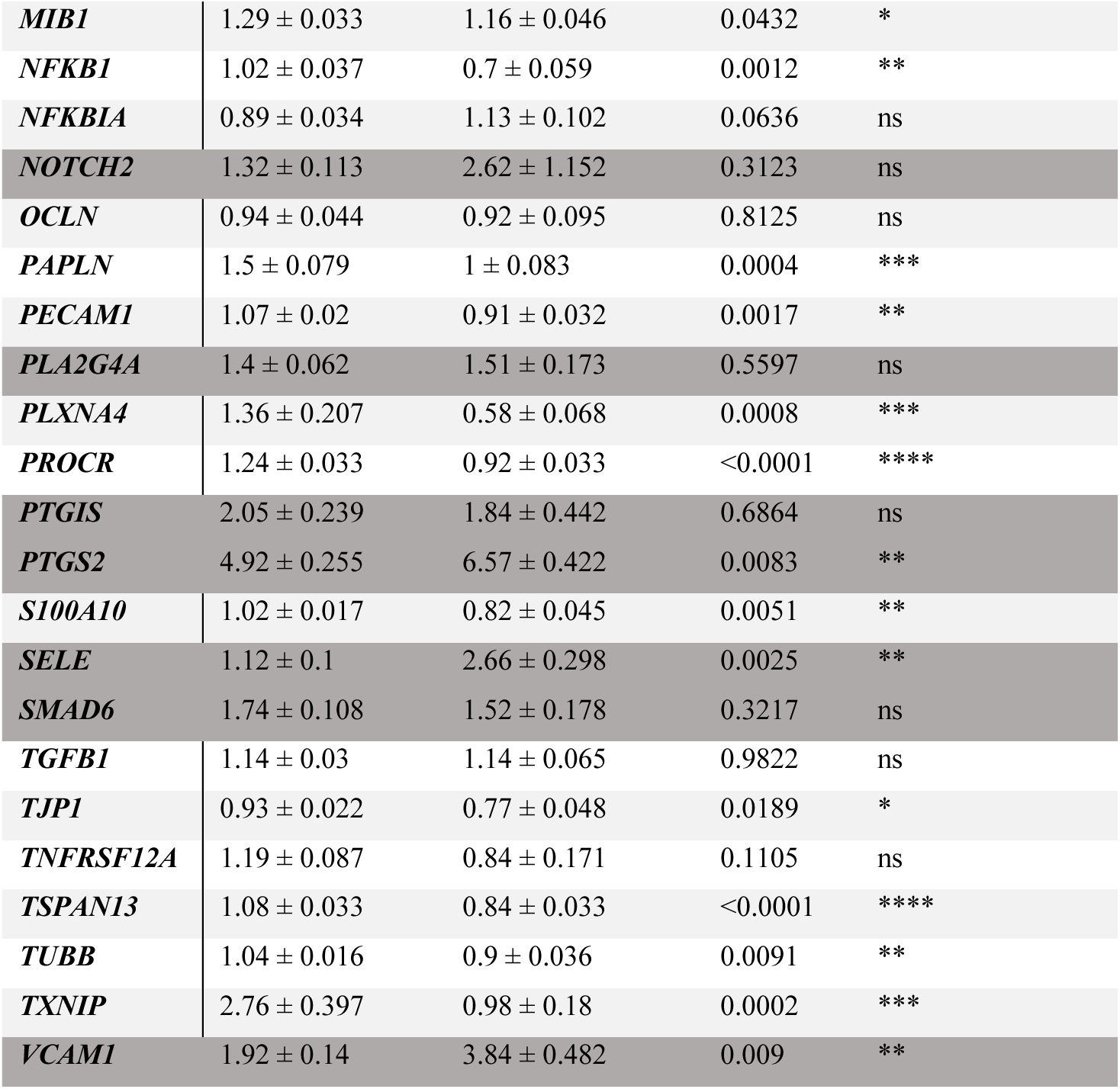
Comparing the relative gene expression of HBMEC co-cultured with either binding IE or non-binding IE for each gene in the panel. Welch’s T-test was performed to compare the mean FC expression of HBMEC incubated with either binding or no-binding IE. All samples were from a 6-hour time point. Data from UM and CM patient-derived parasite isolates were combined to form a group of binding parasites, while two experiments with non-binding SBP1KO parasites were performed. Relative gene expression was calculated using the ΔΔCt method using GAPDH as the endogenous control and the corresponding experimental 6-hour HBMEC-media sample as the reference sample (baseline is FC=1). The mean Fold Change (FC) and standard error of the mean (SEM) for each gene were calculated using all experimental samples and technical replicates (SBP1KO= SBP1-knockout parasites, n = sample size). Rows shaded in dark grey have FC>1.5 for one of the HBMEC groups. P-value summary defined= *,0.01-0.05; **, 0.001-0.01; ***, 0.0001-0.001; ****, <0.0001.

**Supplementary Table 5:**
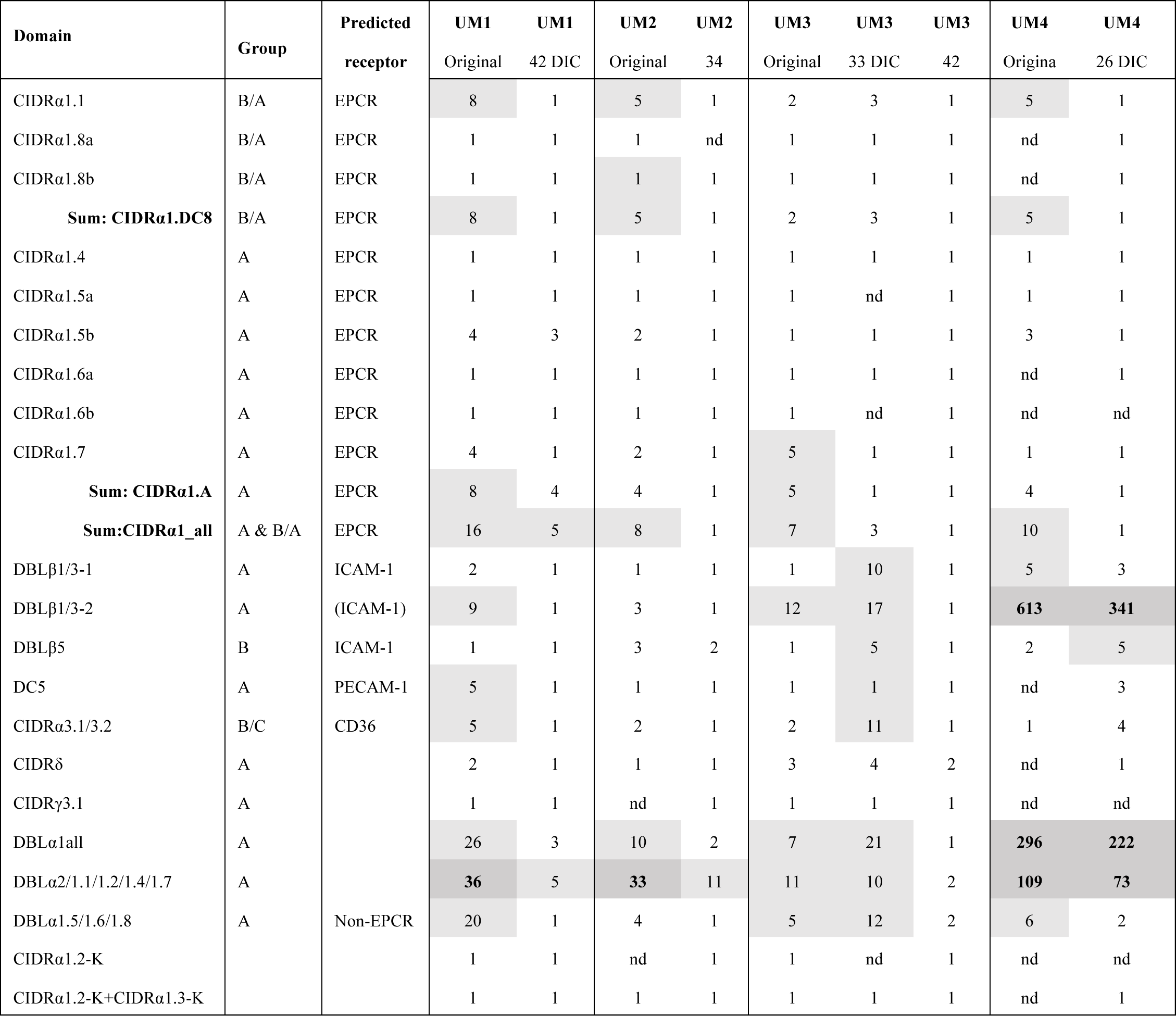
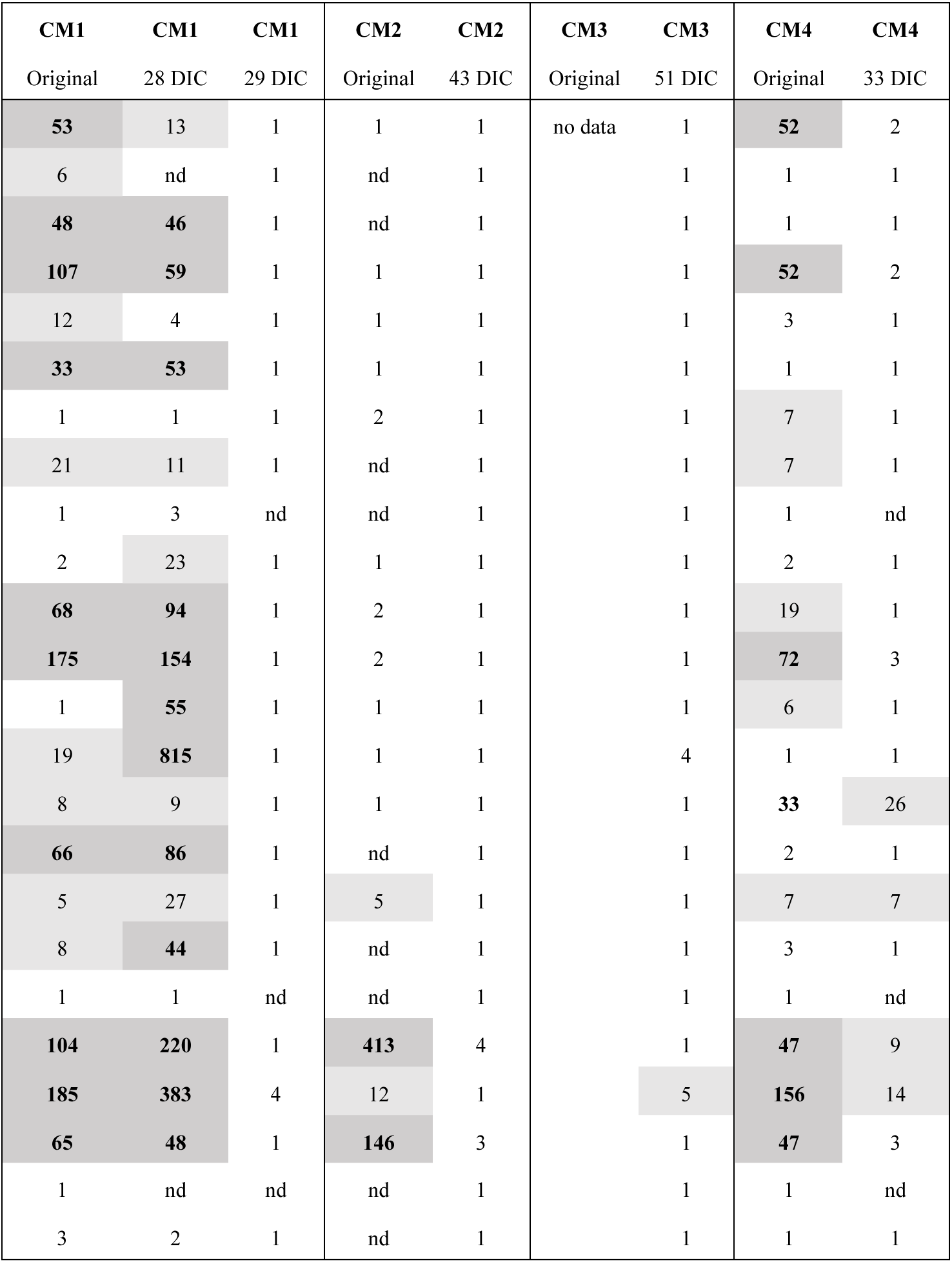
*var* genotyping of patient-derived isolates after culturing by qPCR. The primer sets detect multiple group A and A/B *var* domains and have been used to genotype the patient isolates at the time of isolation (original) (Storm et al, ref 26). For the UM and CM-derived isolates, genotyping was performed after a number of days in culture (DIC), as close to the DIC used for the co-culture experiments. The transcript unit (Tu) was calculated per primer set, relative to the transcripts of the endogenous housekeeping genes *seryl-tRNA synthetase* and *aldolase*, and for the combined (sum) CIDRα1 domains. A Tu value of 1 equals low abundance transcript and a value of 32 equals transcript levels as the endogenous housekeeping genes and values ≥32 are highlighted in bold in the dark grey cells. Tu values ≥5 correspond to moderate transcript levels and are highlighted in the light grey cells. Nd: not determined. Receptors for the PfEMP1 domains are listed with names in brackets as probable and left blank when unknown. For a detailed description of the domains see Mkumbaye et al ^60^.

**Supplementary Table 6:** Determination of prostaglandin endoperoxide synthase 2 (PTGS2) and prostacyclin production in HBMEC-IE co-cultures by ELISA. Prostacyclin has a short half-life and therefore its hydrolysis product, 6-keto prostaglandin F1α (6-keto PGF1α), was detected in culture medium after co-culture with IE or RBC. PTGS2 was detected in HBMEC cell lysate after co-culture and calculated as ng PTGS2 per mg total lysate protein. The mean ± SD of 2 technical replicates is shown and per experiment the ratio of the enzyme concentrations for HBMEC-IE and HBMEC-RBC was calculated. 4 UM-derived isolates, 4 CM-derived isolates and SBP1-KO were co-cultured for 6 hours and for the CM4 isolate and SBP1-KO a co-culture of 20 hours was also performed (experiment 12 and 13). As control, HBMEC were incubated with 10 ng/ml TNF for 6 (experiment C2) and 20 hours (experiment C2 and C3). Thrombin is a fast-acting stimulator of prostacyclin production and the concentration of 6-keto PGF1α was detected in HBMEC after 15- and 30-minutes incubation with 5 nM thrombin (experiment C3). Statistical significance between HBMEC-IE and corresponding HBMEC-RBC and or control condition and medium was determined by unpaired t-test with Welch correction with * p-value < 0.05 and ** p-value < 0.01.

| Exp | Condition | [6ketoPGF1α] pg/ml | | [PTGS2] ng/mg protein | |
| --- | --- | --- | --- | --- | --- |
| | | Mean ± SD | IE/RBC | Mean ± SD | IE/RBC |
| 1 | 6h RBC | 608 ± 43 | | | |
| 1 | 6h UM3-1 | 996 ± 136 | 1.64 | | |
| 1 | 6h UM3-2 | 1046 ± 13 * | 1.72 | | |
| 1 | 6h CM2 | 1153 ± 271 | 1.90 | | |
| 2 | 6h RBC | 1621 ± 390 | | | |
| 2 | 6h UM1 | 952 ± 133 | 0.59 | | |
| 2 | 6h CM3 | 1256 ± 162 | 0.77 | | |
| 3 | 6h RBC | 1679 ± 208 | | | |
| 3 | 6h CM1-1 | 1505 ± 73 | 0.90 |  |  |
| 3 | 6h CM1-2 | 1611 ± 320 | 0.96 |  |  |
| 4 | 6h RBC | 437 ± 38 |  |  |  |
| 4 | 6h UM2 | 803 ± 22 * | 1.84 |  |  |
| 5 | 6h RBC | 664 ± 10 |  | 16.2 ± 0.3 |  |
| 5 | 6h CM3 | 358 ± 100 | 0.54 | 19.1 ± 1.2 | 1.2 |
| 5 | 6h UM1 | 434 ± 56 | 0.65 | 10.8 ± 1.7 | 0.7 |
| 6 | 6h RBC | 772 ± 182 |  | 18.0 ± 1.0 |  |
| 6 | 6h CM4 | 694 ± 105 | 0.90 | 11.4 ± 0.6 * | 0.6 |
| 7 | 6h RBC | 1309 ± 265 |  | 19.0 ± 2.4 |  |
| 7 | 6h UM3 | 1018 ± 79 | 0.78 | 10.0 ± 0.2 | 0.5 |
| 8 | 6h RBC | 1194 ± 46 |  | 21.0 ± 0.6 |  |
| 8 | 6h UM2 | 5720 ± 1378 | 4.79 | 17.4 ± 1.2 | 0.8 |
| 9 | 6h RBC | 396 ± 19 |  | 19.4 ± 1.1 |  |
| 9 | 6h CM1 | 695 ± 12 ** | 1.75 | 13.2 ± 1.3 * | 0.7 |
| 10 | 6h RBC | 1595 ± 114 |  |  |  |
| 10 | 6h UM4 | 2013 ± 244 | 1.26 |  |  |
| 11 | 6h RBC | 4192 ± 1 |  |  |  |
| 11 | 6h SBP1-KO-1 | 7213 ± 561 | 1.72 |  |  |
| 12 | 6h RBC | 1423 ± 109 |  | 13.7 ± 0.1 |  |
| 12 | 6h CM4 | 1871 ± 98 | 1.32 | 12.5 ± 2.4 | 0.9 |
| 12 | 20h RBC | 3754 ± 170 |  | 14.2 ± 1.5 |  |
| 12 | 20h CM4 | 7382 ± 1233 | 1.97 | 9.0 ± 1.4 | 0.6 |
| 13 | 6h RBC | 651 ± 11 |  | 15.3 ± 0.7 |  |
| 13 | 6h SBP1-KO | 480 ± 165 | 0.74 | 14.4 ± 2.1 | 0.9 |
| 13 | 20h RBC | 2729 ± 113 |  | 13.2 ± 1.0 |  |
| 13 | 20h SBP1-KO | 8526 ± 1295 | 3.12 | 11.6 ± 3.1 | 0.9 |
| C1 | 30 min medium | 100 ± 0 |  |  |  |
| C1 | 15 min thrombin (5 nM) | 1071 ± 104 * | 10.74 |  |  |
| C1 | 30 min thrombin (5 nM) | 826 ± 156 | 8.29 |  |  |
| C2 | 6h medium | 264 ± 0 |  |  |  |
| C2 | 6h TNF (10 ng/ml) | 909 ± 62 * | 3.44 |  |  |
| C2 | 20h medium | 685 ± 152 |  |  |  |
| C2 | 20h TNF (10 ng/ml) | 2234 ± 1374 | 3.26 |  |  |
| C3 | 20h medium | 178 ± 13 |  | 123.7 ± 14.9 |  |
| C3 | 20h TNF (10 ng/ml) | 1200 ± 70 * | 6.73 | 118.5 ± 19.4 | 1.0 |

**Supplementary Figure 3:**
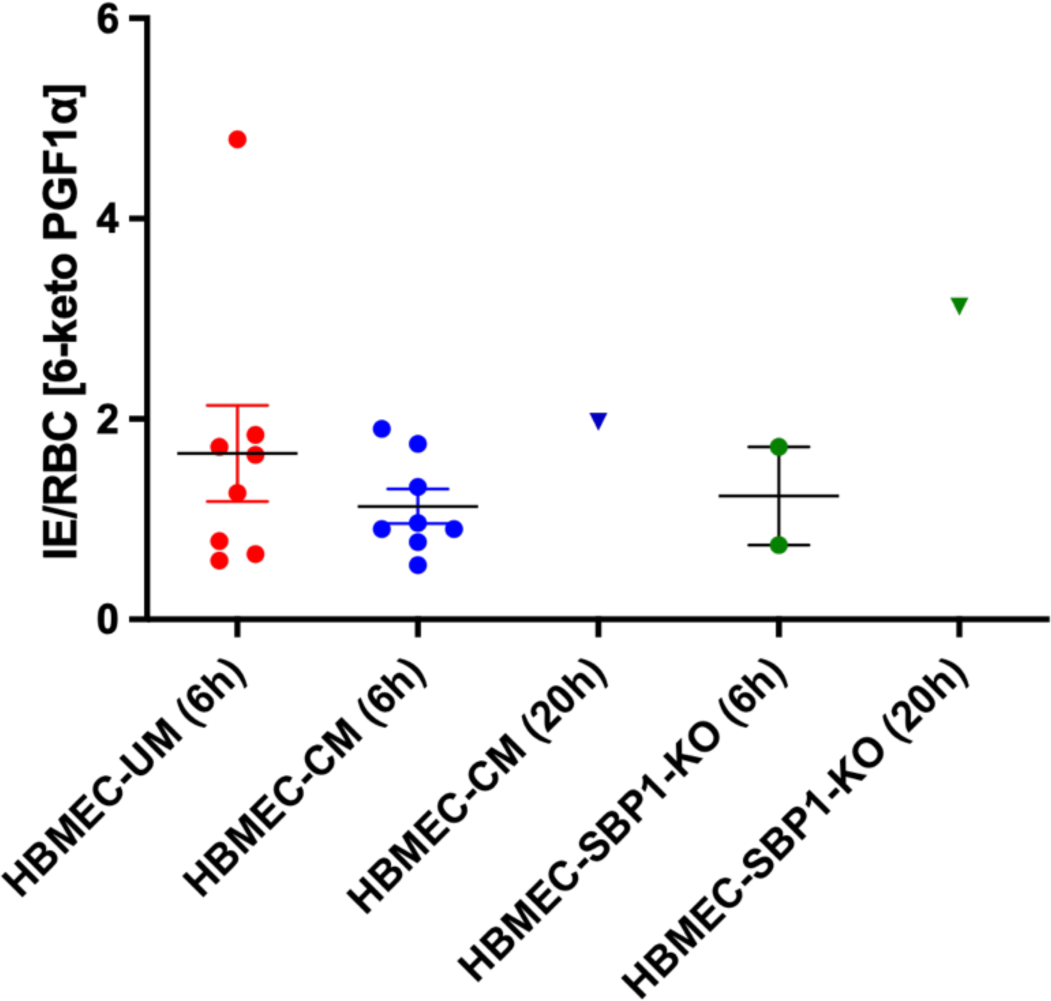
Ratio of 6-keto prostaglandin F1α concentrations in HBMEC-IE and HBMEC-RBC co-cultures. 6-keto PGF1α was detected in culture medium after co-culture with IE or RBC and per experiment the ratio of the enzyme concentrations for HBMEC-IE and HBMEC-RBC was calculated (data in Supplementary Table 4). Plotted are mean ± SD of the ratios for the co-cultures with CM and UM-derived isolates and the SBP1-KO strain.

**Supplementary Table 7:**
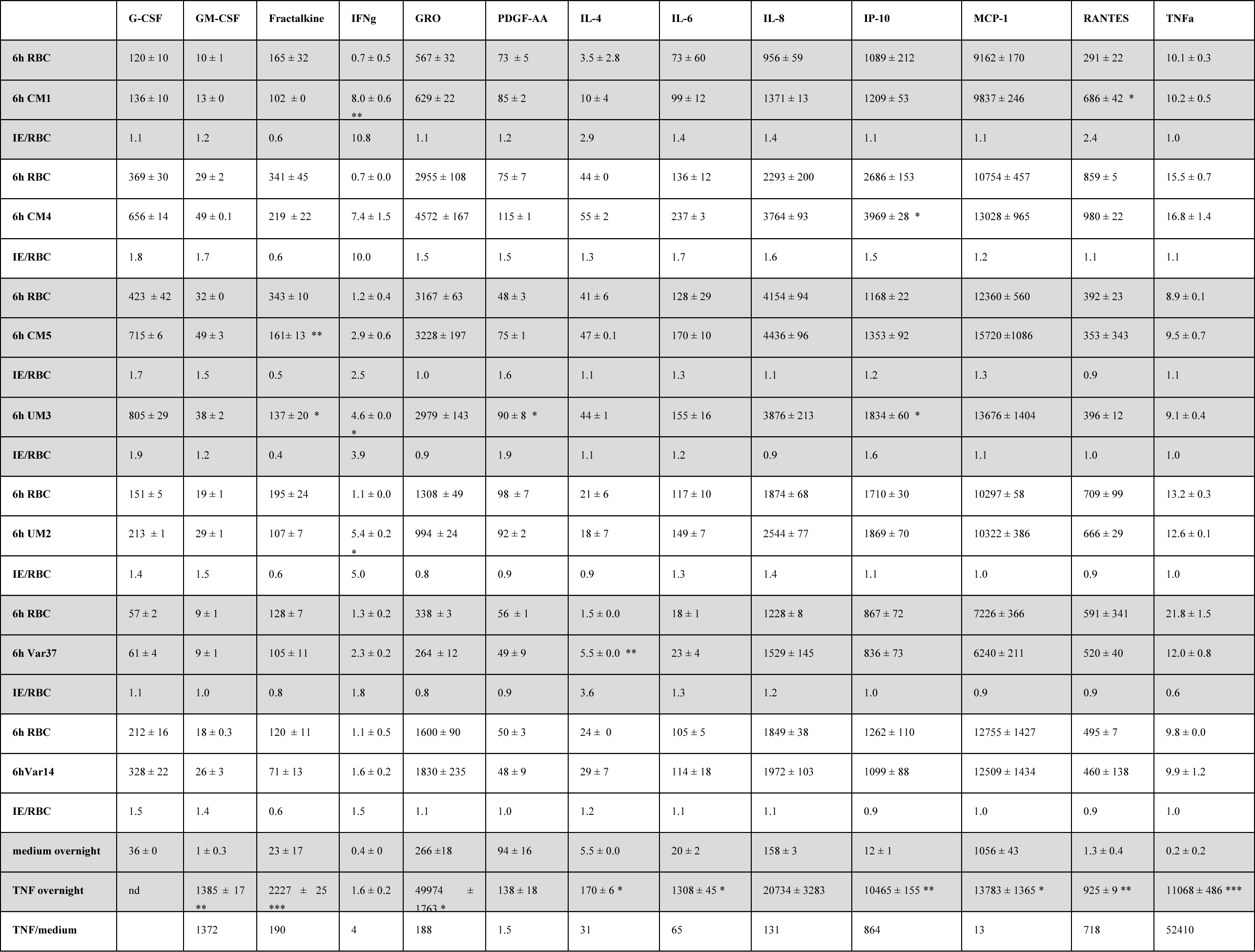
Production of cytokines and chemokines in HBMEC-IE co-cultures. A panel of 41 secreted cytokines and chemokines were measured in the co-culture medium of 6 hours co-culture of HBMEC-IE and HBMEC-RBC by Luminex. The results (pg/ml) are shown for 13 cytokines of which the concentration was higher than 5 pg/ml for 3 CM-derived isolates, 2 UM-derived isolates and 2 lab strains: IT4var14 and IT4var37. A positive control of HBMEC activated by 10 ng/ml TNF for 16 hours was included. The mean ± SD of two technical replicates is shown with significance between HBMEC-IE and HBMEC-RBC calculated by unpaired t-test with Welch correction, with * p-value < 0.05, ** p-value < 0.01 and *** p-value <0. 001.The ratio of the values of HBMEC-IE and HBEC-RBC was calculated per experiment. Note that the CM5 isolate was only included in this multiplex experiment and was not part of all the other experiments described.

**Supplementary Figure 4:**
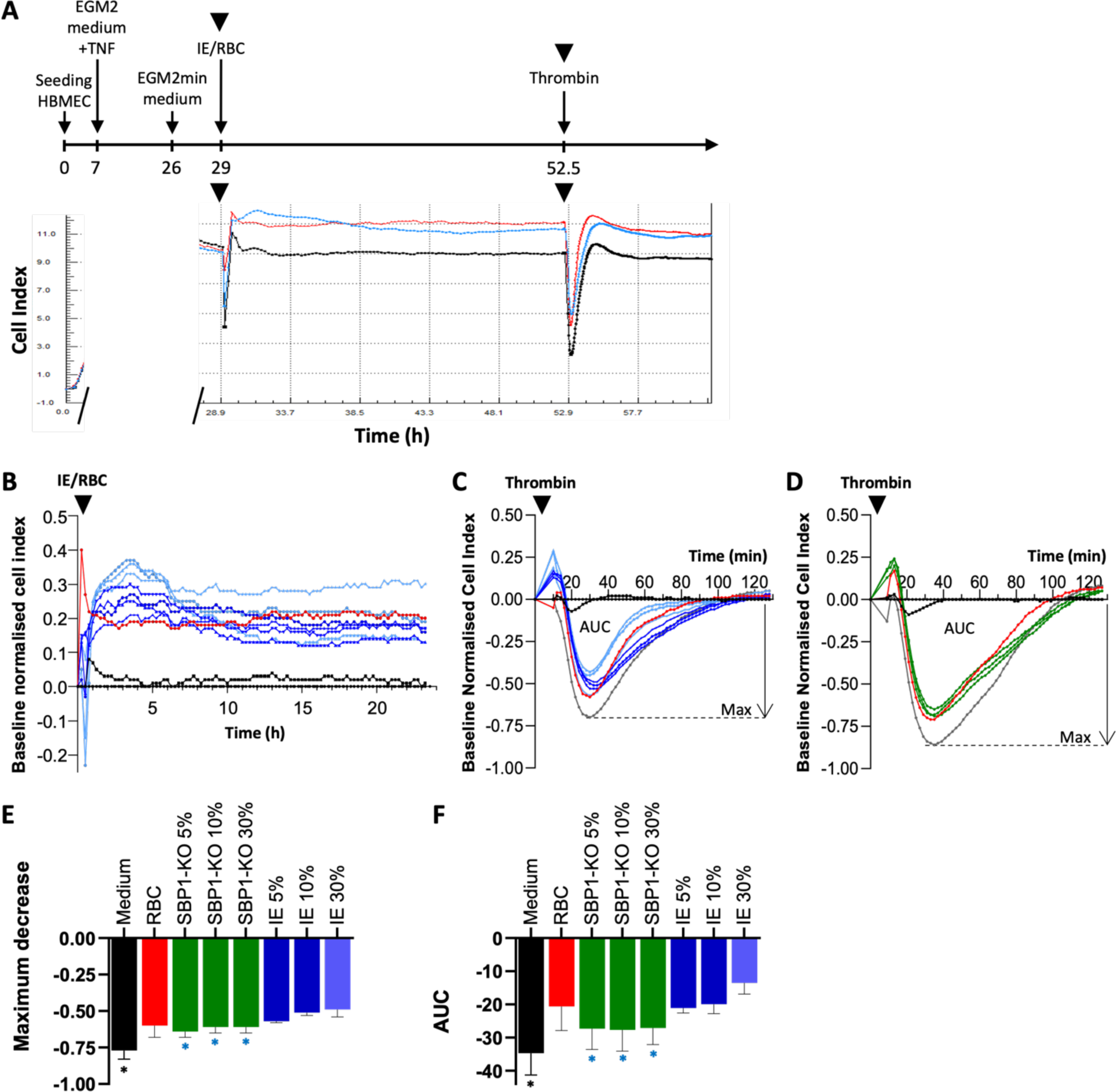
The effect of the patient-derived isolates and SBP1-KO on HBMEC barrier function determined by trans endothelial electrical resistance. (A) Experimental timeline (not to scale) and a representative cell index trace for IE and thrombin induced changes in HBMEC barrier function. HBMEC were seeded in the E-plate and after approximately 7 hrs the cells were stimulated with 10 ng/ml TNF. The next day, medium was replaced with EGM2min medium and after 2-3 hours, IE at 30% parasitaemia (light blue) or RBC (red), both at 1% HCT, were added. The black line represents medium. After approximately 24 hrs, 5 nM thrombin was added from a concentrated stock solution and cell index was monitored for an additional 10 hours. (B) Cell index trace for the effect of different patient isolates and RBC on HBMEC barrier function of a representative experiment. Cell index was normalised at the time point immediately prior to addition of RBC and IE, indicated by the black triangle, and medium only (black line) was set as baseline. This baseline normalised cell index (BNCI) is shown for 2 UM and 2 CM-derived isolates at 10% parasitaemia (dark blue), 1 UM and 2 CM-derived isolates at 30% parasitaemia (light blue), RBC (red) or replicate medium (black). (C) Thrombin induced decrease in barrier function after 24-hour exposure to patient isolates or RBC. Cell index was normalised at the time point immediately prior to addition of 5 nM thrombin, indicated by the black triangle, and medium without thrombin (black line) was set as baseline. BNCI is shown for medium (dark grey) with the maximum response indicated (max), RBC (red), 1 UM and 2 CM-derived isolates at 30% parasitaemia (light blue) and at 10% parasitaemia (dark blue). (D) As in panel C, but with SBP1-KO at 30% and 10% (green). (E) Maximum decrease of BNCI by thrombin was determined for all co-culture conditions and mean ± SD are depicted for: control medium (n = 4), RBC (n = 4), SBP1-KO at 5% (n = 3), 10% (n = 3) and 30% parasitaemia (n = 3) and combined patient isolates (IE) at 5% (n = 2), 10% (n = 5) and 30% parasitaemia (n = 5). (F) Area under the curve (AUC) after thrombin induced changes in BNCI was calculated and mean ± SD are depicted for the conditions listed in E. Statistical significance between the various conditions compared to RBC was calculated by unpaired t-test with Welch correction, * p-value < 0.05 (black asterisk). Significance between 30% IE and SBP1-KO was also calculated (blue asterisk).

## Supplementary Methods

### *var* Genotyping of Patient-Derived Isolates after Culturing

Genotyping of the isolates after culturing was performed by qPCR using selected primers from a *var* gene primer panel as described by Storm et al ^26^, with details of the primer design and description of the var domains described by Mkumbaye et al ^60^. RNA was isolated from TRIZOL-stored ring stage IE, DNase treated (TURBO™ DNase, Ambion), cDNA synthesised (Tetro cDNA Synthesis Kit, Bioline) and qPCR carried out with SYBR Green PCR Master Mix (QuantiTect, Qiagen). The endogenous housekeeping genes *seryl-tRNA synthetase* and *aldolase* were used to determine the relative levels of *var* transcripts using the formulae ΔCt *var*-primer = Ct *var* primer minus Ct average of endogenous primers. The transcript unit (Tu) was then calculated as Tu = 2^(5-ΔCt^ ^var-primer)^. Any low abundance transcripts with a ΔCt *var*-primer >5 was assigned a value of 5, thus subsequently a Tu value of 1. A Tu value of 32 equates to equal transcript levels as the endogenous control genes ^60^. The Tu of the patient isolates at the time of isolation from peripheral blood was determined previously ^26^.

### Preparation of HBMEC-IE Co-culture Samples for Phenotypic Analysis

After 6 or 20 hour co-culture with patient isolates, SBP1-KO, RBC or additional controls, co-culture medium was collected, centrifuged for 5 min at 900 g to remove IE, RBC or cellular debris and the supernatant frozen at −80 °C. For cell lysates, HBMEC were washed twice with warm EGM2 medium without supplements and lysed in 50 µl lysis buffer (RIPA buffer (Sigma, R0278) with HALT™ protease inhibitor cocktail (Thermo Scientific, 78429) for 10 minutes on ice. The lysed cells were transferred from the well into a tube, placed on ice for an additional 25 minutes, centrifuged for 5 min at 13,000 g at 4 °C and the supernatant frozen at −80 °C.

### Determination of the Concentrations of Prostacyclin and Prostaglandin Endoperoxide Synthase 2

Prostacyclin has a short half-life and therefore its hydrolysis product, 6-keto prostaglandin F1α (6-keto PGF1α), was detected in co-culture medium by competitive ELISA according to the manufacturer’s instructions (Cayman Chemicals, 515211). The four UM and four CM-derived isolates and SBP1-KO were co-cultured for 6 hours with their respective RBC as control. The CM4 isolate and SBP1-KO were also co-cultured for 20 hours. As additional controls, HBMEC were incubated with 10 ng/ml TNF for 6 and 20 hours and with 5 nM thrombin for 15 and 30 minutes. Co-culture medium of two independent 6 hour co-culture experiments with the patient isolates, except UM4 and CM2, and SBP1-KO were used, and all samples were measured in duplicate using 2 ELISA plates on the same day. All samples were diluted 2 times, but the 20 hour TNF stimulation sample was diluted 5 times. The concentration of 6-keto PGF1α was calculated using the provided standards.

Prostaglandin endoperoxide synthase 2 (PTGS2) was detected in HBMEC cell lysate by ELISA according to the manufacturer’s instructions (Reddot, RD-PTGS2-Hu). Lysates were diluted 5 times and for duplicate samples the concentration of PTGS2 was calculated using the provided standards. On the same day, total protein concentration of the lysates was determined with the BCA kit (Biorad, 5000002) with BSA as standard and ng PTGS2 per mg total protein calculated.

### Detection of Cytokines and Chemokines

A panel of 41 secreted cytokines and chemokines was measured in the co-culture medium of 6 hours co-culture of HBMEC-IE, HBMEC-RBC and controls using the human cytokine/chemokine 41 plex Immunology Multiplex Assay according to the manufacturer’s instructions (Merck, HCYTMAG-60K-PX41). In addition to three CM and two UM-derived isolates, two lab strains, IT4var14 and IT4var37, and a positive control of HBMEC activated by 10 ng/ml TNF for 16 hours were included. The panel included: sCD40L, Eotaxin, FLT-3L, Fractalkine, G-CSF, GM-CSF, GROα, IFNα2, IFNγ, IL-1α, IL-1β, IL-1RA, IL-2, IL-3, IL-4, IL-5, IL-6, IL-7, IL-8, IL-9, IL-10, IL-12 (p40), IL-12 (p70), IL-13, IL-15, IL-17A, IL-17E/IL-25, IL-17F, IL-18, IL-22, IL-27, IP-10, MCP-1, MCP-3, M-CSF, MDC, MIG, MIP-1α, MIP-1β, PDGF-AA, PDGF-AB/BB, RANTES, TGFα, TNFα, TNFβ, VEGF-A.

### Measuring HBMEC Barrier Integrity

Barrier function was measured by real time Trans Endothelial Electrical Resistance (TEER) analysis with the xCELLigence® RTCA S16 system (ACEA Biosciences). HBMEC cells were seeded at 50,000 cells/cm^2^ (1×10^4^ cells) in Attachment Factor coated E-Plates 16 PET (ACEA Biosciences) in EGM2 medium and the arbitrary cell index (CI) was recorded. After approximately 7 hrs the medium was replaced with EGM2 medium with 10 ng/ml TNF and the next day, medium was replaced with EGM2min medium. IE suspensions at 30%, 10% or 5% parasitaemia were prepared at 1% HCT in EGM2min medium and the medium in the E-plate replaced with 100 µl suspension, equivalent to 3×106, 1×106 and 0.5×106 IE, respectively. RBC at 1% HCT and medium only were used as control.

After approximately 24 hrs, 5 nM thrombin was added from a concentrated stock solution and cell index was monitored for an additional 10 hours, starting at 2 minute intervals. For analysis, the cell index was normalised either at the time point immediately prior to addition of IE or thrombin (normalised cell index) and medium only was set as baseline (baseline normalised cell index, BNCI). To determine the effect of thrombin, the maximum decrease in BNCI and the area under the curve after recovery to baseline was calculated.

## Notes

### Competing Interest Statement

The authors have declared no competing interest.

### Summary of Updates

This version of the manuscript has been revised following reviewer comments. Figures and tables have been put into supplementary, and one figure has been removed, along with other revisions.

## References

1 World Malaria Report 2023. Geneva: World Health Organization; 2023. Licence: CC BY-NC-SA 3.0 IGO. (2023).

2 Phillips, M. A. et al. Malaria. Nat Rev Dis Primers 3, 17050, doi:10.1038/nrdp.2017.50 (2017).

3 Kojom Foko, L. P., Kumar, A., Hawadak, J. & Singh, V. Plasmodium cynomolgi in humans: current knowledge and future directions of an emerging zoonotic malaria parasite. Infection 51, 623–640, doi:10.1007/s15010-022-01952-2 (2023).

4 de Oliveira, T. C. et al. Plasmodium simium: Population Genomics Reveals the Origin of a Reverse Zoonosis. J Infect Dis 224, 1950–1961, doi:10.1093/infdis/jiab214 (2021).

5 Song, X. et al. Cerebral malaria induced by plasmodium falciparum: clinical features, pathogenesis, diagnosis, and treatment. Front Cell Infect Microbiol 12, 939532, doi:10.3389/fcimb.2022.939532 (2022).

6 Seydel, K. B. et al. Brain swelling and death in children with cerebral malaria. N Engl J Med 372, 1126–1137, doi:10.1056/NEJMoa1400116 (2015).

7 Mohanty, S. et al. Magnetic Resonance Imaging of Cerebral Malaria Patients Reveals Distinct Pathogenetic Processes in Different Parts of the Brain. mSphere 2, doi:10.1128/mSphere.00193-17 (2017).

8 Storm, J. & Craig, A. G. Pathogenesis of cerebral malaria--inflammation and cytoadherence. Front Cell Infect Microbiol 4, 100, doi:10.3389/fcimb.2014.00100 (2014).

9 Ramachandran, A. & Sharma, A. Dissecting the mechanisms of pathogenesis in cerebral malaria. PLoS Pathog 18, e1010919, doi:10.1371/journal.ppat.1010919 (2022).

10 Pasternak, N. D. & Dzikowski, R. PfEMP1: an antigen that plays a key role in the pathogenicity and immune evasion of the malaria parasite Plasmodium falciparum. Int J Biochem Cell Biol 41, 1463–1466, doi:10.1016/j.biocel.2008.12.012 (2009).

11 Smith, J. D. The role of PfEMP1 adhesion domain classification in Plasmodium falciparum pathogenesis research. Mol Biochem Parasitol 195, 82–87, doi:10.1016/j.molbiopara.2014.07.006 (2014).

12 Albrecht-Schgoer, K., Lackner, P., Schmutzhard, E. & Baier, G. Cerebral Malaria: Current Clinical and Immunological Aspects. Front Immunol 13, 863568, doi:10.3389/fimmu.2022.863568 (2022).

13 Kraemer, S. M. & Smith, J. D. A family affair: var genes, PfEMP1 binding, and malaria disease. Curr Opin Microbiol 9, 374–380, doi:10.1016/j.mib.2006.06.006 (2006).

14 Bernabeu, M. & Smith, J. D. EPCR and Malaria Severity: The Center of a Perfect Storm. Trends Parasitol 33, 295–308, doi:10.1016/j.pt.2016.11.004 (2017).

15 Kessler, A. et al. Linking EPCR-Binding PfEMP1 to Brain Swelling in Pediatric Cerebral Malaria. Cell Host Microbe 22, 601–614 e605, doi:10.1016/j.chom.2017.09.009 (2017).

16 Turner, L. et al. Severe malaria is associated with parasite binding to endothelial protein C receptor. Nature 498, 502–505, doi:10.1038/nature12216 (2013).

17 Moxon, C. A., Gibbins, M. P., McGuinness, D., Milner, D. A., Jr. & Marti, M. New Insights into Malaria Pathogenesis. Annu Rev Pathol 15, 315–343, doi:10.1146/annurev-pathmechdis-012419-032640 (2020).

18 Lennartz, F. et al. Structure-Guided Identification of a Family of Dual Receptor-Binding PfEMP1 that Is Associated with Cerebral Malaria. Cell Host Microbe 21, 403–414, doi:10.1016/j.chom.2017.02.009 (2017).

19 Avril, M., Bernabeu, M., Benjamin, M., Brazier, A. J. & Smith, J. D. Interaction between Endothelial Protein C Receptor and Intercellular Adhesion Molecule 1 to Mediate Binding of Plasmodium falciparum-Infected Erythrocytes to Endothelial Cells. mBio 7, doi:10.1128/mBio.00615-16 (2016).

20 Gallego-Delgado, J. & Rodriguez, A. Rupture and Release: A Role for Soluble Erythrocyte Content in the Pathology of Cerebral Malaria. Trends Parasitol 33, 832–835, doi:10.1016/j.pt.2017.06.005 (2017).

21 Adams, Y. et al. Plasmodium falciparum erythrocyte membrane protein 1 variants induce cell swelling and disrupt the blood-brain barrier in cerebral malaria. J Exp Med 218, doi:10.1084/jem.20201266 (2021).

22 Bernabeu, M. et al. Binding Heterogeneity of Plasmodium falciparum to Engineered 3D Brain Microvessels Is Mediated by EPCR and ICAM-1. mBio 10, doi:10.1128/mBio.00420-19 (2019).

23 Zuniga, M. et al. Plasmodium falciparum and TNF-alpha Differentially Regulate Inflammatory and Barrier Integrity Pathways in Human Brain Endothelial Cells. mBio 13, e0174622, doi:10.1128/mbio.01746-22 (2022).

24 Azasi, Y. et al. Infected erythrocytes expressing DC13 PfEMP1 differ from recombinant proteins in EPCR-binding function. Proc Natl Acad Sci U S A 115, 1063–1068, doi:10.1073/pnas.1712879115 (2018).

25 Othman, B. et al. Different PfEMP1-expressing Plasmodium falciparum variants induce divergent endothelial transcriptional responses during co-culture. PLoS One 18, e0295053, doi:10.1371/journal.pone.0295053 (2023).

26 Storm, J. et al. Cerebral malaria is associated with differential cytoadherence to brain endothelial cells. EMBO Mol Med 11, doi:10.15252/emmm.201809164 (2019).

27 Maier, A. G. et al. Skeleton-binding protein 1 functions at the parasitophorous vacuole membrane to traffic PfEMP1 to the Plasmodium falciparum-infected erythrocyte surface. Blood 109, 1289–1297, doi:10.1182/blood-2006-08-043364 (2007).

28 Livak, K. J. & Schmittgen, T. D. Analysis of relative gene expression data using real-time quantitative PCR and the 2(-Delta Delta C(T)) Method. Methods 25, 402–408, doi:10.1006/meth.2001.1262 (2001).

29 Madkhali, A. M. et al. An analysis of the binding characteristics of a panel of recently selected ICAM-1 binding Plasmodium falciparum patient isolates. PLoS One 9, e111518, doi:10.1371/journal.pone.0111518 (2014).

30 Howard, C., Joof, F., Hu, R., Smith, J. D. & Zheng, Y. Probing cerebral malaria inflammation in 3D human brain microvessels. Cell Rep 42, 113253, doi:10.1016/j.celrep.2023.113253 (2023).

31 Karsan, A., Yee, E., Kaushansky, K. & Harlan, J. M. Cloning of human Bcl-2 homologue: inflammatory cytokines induce human A1 in cultured endothelial cells. Blood 87, 3089–3096 (1996).

32 Peters, J. M., Fowler, E. V., Krause, D. R., Cheng, Q. & Gatton, M. L. Differential changes in Plasmodium falciparum var transcription during adaptation to culture. J Infect Dis 195, 748–755, doi:10.1086/511436 (2007).

33 Mkumbaye, S. I. et al. Cellulose filtration of blood from malaria patients for improving ex vivo growth of Plasmodium falciparum parasites. Malar J 16, 69, doi:10.1186/s12936-017-1714-2 (2017).

34 Cooke, B. M. et al. A Maurer’s cleft-associated protein is essential for expression of the major malaria virulence antigen on the surface of infected red blood cells. J Cell Biol 172, 899–908, doi:10.1083/jcb.200509122 (2006).

35 Hadjilaou, A., Brandi, J., Riehn, M., Friese, M. A. & Jacobs, T. Pathogenetic mechanisms and treatment targets in cerebral malaria. Nat Rev Neurol 19, 688–709, doi:10.1038/s41582-023-00881-4 (2023).

36 Moxon, C. A. et al. Parasite histones are toxic to brain endothelium and link blood barrier breakdown and thrombosis in cerebral malaria. Blood Adv 4, 2851–2864, doi:10.1182/bloodadvances.2019001258 (2020).

37 Bachmann, A. et al. Highly co-ordinated var gene expression and switching in clinical Plasmodium falciparum isolates from non-immune malaria patients. Cell Microbiol 13, 1397–1409, doi:10.1111/j.1462-5822.2011.01629.x (2011).

38 Lee, W. C., Russell, B. & Renia, L. Sticking for a Cause: The Falciparum Malaria Parasites Cytoadherence Paradigm. Front Immunol 10, 1444, doi:10.3389/fimmu.2019.01444 (2019).

39 Chan, J. A. et al. A single point in protein trafficking by Plasmodium falciparum determines the expression of major antigens on the surface of infected erythrocytes targeted by human antibodies. Cell Mol Life Sci 73, 4141–4158, doi:10.1007/s00018-016-2267-1 (2016).

40 Possemiers, H. et al. Skeleton binding protein-1-mediated parasite sequestration inhibits spontaneous resolution of malaria-associated acute respiratory distress syndrome. PLoS Pathog 17, e1010114, doi:10.1371/journal.ppat.1010114 (2021).

41 Watson, E. C., Grant, Z. L. & Coultas, L. Endothelial cell apoptosis in angiogenesis and vessel regression. Cell Mol Life Sci 74, 4387–4403, doi:10.1007/s00018-017-2577-y (2017).

42 Vogler, M. BCL2A1: the underdog in the BCL2 family. Cell Death Differ 19, 67–74, doi:10.1038/cdd.2011.158 (2012).

43 Noble, K. E., Wickremasinghe, R. G., DeCornet, C., Panayiotidis, P. & Yong, K. L. Monocytes stimulate expression of the Bcl-2 family member, A1, in endothelial cells and confer protection against apoptosis. J Immunol 162, 1376–1383 (1999).

44 Xu, S. et al. The zinc finger transcription factor, KLF2, protects against COVID-19 associated endothelial dysfunction. Signal Transduct Target Ther 6, 266, doi:10.1038/s41392-021-00690-5 (2021).

45 SenBanerjee, S. et al. KLF2 Is a novel transcriptional regulator of endothelial proinflammatory activation. J Exp Med 199, 1305–1315, doi:10.1084/jem.20031132 (2004).

46 Hamik, A. et al. Kruppel-like factor 4 regulates endothelial inflammation. J Biol Chem 282, 13769–13779, doi:10.1074/jbc.M700078200 (2007).

47 Sangwung, P. et al. KLF2 and KLF4 control endothelial identity and vascular integrity. JCI Insight 2, e91700, doi:10.1172/jci.insight.91700 (2017).

48 Zhou, G. et al. Endothelial Kruppel-like factor 4 protects against atherothrombosis in mice. J Clin Invest 122, 4727–4731, doi:10.1172/JCI66056 (2012).

49 Atkins, G. B. & Jain, M. K. Role of Kruppel-like transcription factors in endothelial biology. Circ Res 100, 1686–1695, doi:10.1161/01.RES.0000267856.00713.0a (2007).

50 Lin, Z. et al. Kruppel-like factor 2 (KLF2) regulates endothelial thrombotic function. Circ Res 96, e48–57, doi:10.1161/01.RES.0000159707.05637.a1 (2005).

51 Ghaleb, A. M. & Yang, V. W. Kruppel-like factor 4 (KLF4): What we currently know. Gene 611, 27–37, doi:10.1016/j.gene.2017.02.025 (2017).

52 Stitham, J., Midgett, C., Martin, K. A. & Hwa, J. Prostacyclin: an inflammatory paradox. Front Pharmacol 2, 24, doi:10.3389/fphar.2011.00024 (2011).

53 Dorris, S. L. & Peebles, R. S., Jr. PGI2 as a regulator of inflammatory diseases. Mediators Inflamm 2012, 926968, doi:10.1155/2012/926968 (2012).

54 Ferguson, C. S. & Tyndale, R. F. Cytochrome P450 enzymes in the brain: emerging evidence of biological significance. Trends Pharmacol Sci 32, 708–714, doi:10.1016/j.tips.2011.08.005 (2011).

55 Avril, M., Benjamin, M., Dols, M. M. & Smith, J. D. Interplay of Plasmodium falciparum and thrombin in brain endothelial barrier disruption. Sci Rep 9, 13142, doi:10.1038/s41598-019-49530-1 (2019).

56 O’Carroll, S. J. et al. Pro-inflammatory TNFalpha and IL-1beta differentially regulate the inflammatory phenotype of brain microvascular endothelial cells. J Neuroinflammation 12, 131, doi:10.1186/s12974-015-0346-0 (2015).

57 Storm, J., Wu, Y., Davies, J., Moxon, C. A. & Craig, A. G. Testing the effect of PAR1 inhibitors on Plasmodium falciparum-induced loss of endothelial cell barrier function. Wellcome Open Res 5, 34, doi:10.12688/wellcomeopenres.15602.3 (2020).

58 Alberelli, M. A. & De Candia, E. Functional role of protease activated receptors in vascular biology. Vascul Pharmacol 62, 72–81, doi:10.1016/j.vph.2014.06.001 (2014).

59 Moxon, C. A. et al. Persistent endothelial activation and inflammation after Plasmodium falciparum Infection in Malawian children. J Infect Dis 209, 610–615, doi:10.1093/infdis/jit419 (2014).

60 Mkumbaye, S. I. et al. The Severity of Plasmodium falciparum Infection Is Associated with Transcript Levels of var Genes Encoding Endothelial Protein C Receptor-Binding P. falciparum Erythrocyte Membrane Protein 1. Infect Immun 85, doi:10.1128/IAI.00841-16 (2017).

